# ACE2-independent SARS-CoV-2 infection and mouse adaption emerge after passage in cells expressing human and mouse ACE2

**DOI:** 10.1101/2021.12.16.473063

**Authors:** Kexin Yan, Troy Dumenil, Bing Tang, Thuy T. Le, Cameron Bishop, Andreas Suhrbier, Daniel J. Rawle

## Abstract

Human ACE2 (hACE2) is the key cell attachment and entry receptor for SARS-CoV-2, with the original SARS-CoV-2 isolates unable to use mouse ACE2 (mACE2). Herein we describe a new system for generating mouse-adapted SARS-CoV-2 *in vitro* by serial passaging virus in co-cultures of cell lines expressing hACE2 and mACE2. Mouse-adapted viruses emerged with up to five amino acid changes in the spike protein, all of which have been seen in human isolates. Mouse-adapted viruses replicated to high titers in C57BL/6J mouse lungs and nasal turbinates, and caused severe lung histopathology. One mouse-adapted virus was also able to replicate efficiently in ACE2-negative cell lines, with ACE2-independent entry by SARS-CoV-2 representing a new biology for SARS-CoV-2 that has potential widespread implications for disease and intervention development.

## INTRODUCTION

Severe acute respiratory syndrome coronavirus 2 (SARS-CoV-2) emerged in 2019, causing a global pandemic of coronavirus disease 2019 (COVID-19)^1^. The human ACE2 (hACE2) receptor is generally held to be the key cell attachment and entry receptor for SARS-CoV-2, with the receptor binding domain (RBD) of the spike protein binding to hACE2^2^. ACE2 binding has deep ancestral origins within the sarbecovirus lineage of coronaviruses^3^, with SARS-CoV, SARS-CoV-2, and SARS-related bat coronaviruses all using ACE2 as their entry receptor^4^. Genetically diverse spike proteins are able to bind ACE2^3,4^, with all the SARS-CoV-2 variants of concern (Alpha, Beta, Gamma, Delta and Omicron) reported to require ACE2 binding for efficient infection^5,6^.

The original isolates of SARS-CoV-2 (Wuhan strain) are unable to bind mouse ACE2 (mACE2) for infection^2^, therefore, to study these viruses a series of mouse models were developed that express hACE2^2,7-10^. Some subsequent SARS-CoV-2 variants of concern emerged to be able to use mACE2, including Alpha, Beta, Gamma, and Omicron, but not Delta^6,11^. In addition, mouse-adapted SARS-CoV-2 have been generated by serial passage of original SARS-CoV-2 isolates in mouse lungs^12-16^ or by reverse genetics^17^ to introduce amino acid changes that allow binding to mACE2.

As infection of wild animals by SARS-CoV-2 is increasingly being reported^18-23^, with omicron also potentially arising from rodents^24,25^, we further sought to characterize the process of mouse adaptation. We have previously reported the use of HEK293T cells that express hACE2 (HEK293T-hACE2) or mACE2 (HEK293T-mACE2) by virtue of lentiviral transduction, with the former, but not the latter, able to support efficient replication of an original SARS-CoV-2 isolate, hCoV-19/Australia/QLD02/2020 (SARS-CoV-2_QLD02_)^2^. We also showed that transduction with mACE2 containing the N31K and H353K mouse-to-human amino acid changes (to generate HEK293T-mACE2^N31K/H353K^ cells), represented the minimum changes required to support SARS-CoV-2 _QLD02_ replication^2^. Herein we serially passaged SARS-CoV-2_QLD02_ in HEK293T-hACE2 or HEK293T-mACE2^N31K/H353K^ cells co-cultured with HEK293T-mACE2 cells, followed by passaging in HEK293T-mACE2 cells to generate five different mouse-adapted (MA) viruses. These MA viruses were able to replicate in both HEK293T-hACE2 and HEK293T-mACE2 cells. These viruses were also able to replicate in C57BL/6J mice, leading to characteristic COVID-19 histopathological lesions and inflammatory responses. Remarkably, one of these viruses (MA1) was also able to replicate in HEK293T cells in the absence of lentiviral ACE2 expression. Furthermore, MA1 replicated in a number of other cell lines that do not express significant levels of ACE2 and do not support SARS-CoV-2_QLD02_ infection. This represents just the second report of productive infection in unmodified ACE2-negative cell lines^26^, with potential implications for the biology of SARS-CoV-2 infections, evolution, tropism, disease and perhaps also intervention development.

## RESULTS

### SARS-CoV-2 _QLD02_ co-culture passaging to select for MA viruses

To investigate the process of mouse adaptation, SARS-CoV-2_QLD02_ was passaged in co-cultures of HEK293T-hACE2 and HEK293T-mACE2, or HEK293T-mACE2^N31K/H353K^ and HEK293T-mACE2 cells, followed by passage in HEK293T-mACE2 cells (Fig. 1a. After 4 co-culture passages, no viruses emerged that were able to replicate efficiently in HEK293T-mACE2 cells (Fig. 1b, Passage 4). However, after 9 co-culture passages, 5 viruses (MA1-5) demonstrated a clear ability to replicate in HEK293T-mACE2 cells (Fig. 1b, Passage 9). As expected, SARS-CoV-2_QLD02_ did not replicate in HEK293T-mACE2 cells (Fig. 1b). All 5 MA viruses showed overt cytopathic effects (CPE) in HEK293T-mACE2 cells 2 days after infection (Fig. 1c).

**Figure 1.**
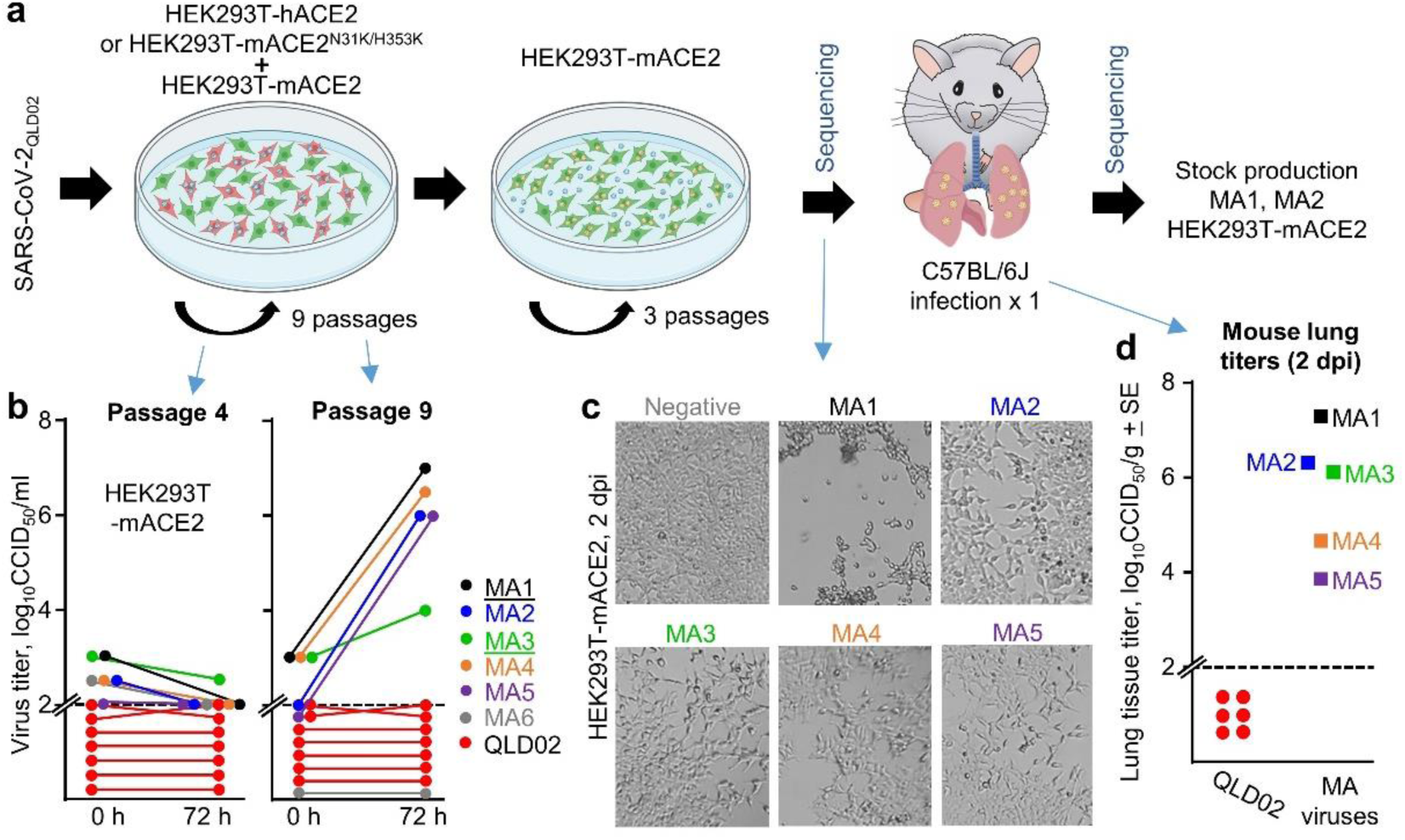
*In vitro* adaptation of SARS-CoV-2_QLD02_ to mACE2 utilization. **a** Schematic of SARS-CoV-2_QLD02_ passaging in HEK293T-hACE2 or HEK293T-mACE2^N31K/H353K^ cells, co-cultured with HEK293T-mACE2 cells (9 passages), followed by passaging in HEK293T-mACE2 cells (3 passages) and mouse infections. Viruses used to infect mice and viruses derived from mice lungs (2 dpi) were sequenced. Stock virus was prepared for MA1 and MA2 from mouse lungs 2 dpi for use in subsequent experiments. **b** Supernatants from passage 4 and 9 (from the co-cultures – blue arrows) were used to infect HEK93T-mACE2 cells and viral growth over 72 hours determined by CCID_50_ assays. Dotted line – limit of detection. MA1 and MA3 (underlined) were derived from HEK293T-hACE2/HEK293T-mACE2 co-cultures, MA2, 4, 5 and 6 from HEK293T-mACE2^N31K/H353K^/HEK293T-mACE2 co-cultures. **c** Inverted light microscopy images of CPE in HEK293T-mACE2 cells infected with passage 9 supernatants at 72 hours post-infection. Images are representative of at least three replicates. **d** MA viruses obtained after 3 passages in HEK293T-mACE2 cells were used to infect C57BL/6J mice. Lung tissue titers are shown for 2 dpi for MA1-5 (n=1 for each) and for SARS-CoV-2_QLD02_ (n=6).

Importantly, these viruses were also able to replicate in lungs of wild-type C57BL/6J mice after intrapulmonary inoculation (via the intranasal route) with 1.5×10^4^ CCID_50_ per mouse (Fig. 1d). MA1 and MA2 showed the highest lung titers, replicating to above 10^6^ CCID_50_/g (Fig. 1d). As expected SARS-CoV-2_QLD02_ was unable to replicate in C57BL/6J mice (Fig. 1d). The co-culture and *in vitro* passaging of SARS-CoV-2_QLD02_ thus generated five viruses able to replicate in wild-type mice.

### Amino acid changes in MA viruses have previously been reported in human isolates

MA1-5 viruses were sequenced before and after infection of C57BL/6J mice (Fig. 1a, Sequencing) by RNA-Seq. Sequences for MA viruses before and after infection of C57BL/6J mice (Fig. 1a, Sequencing) were almost identical (Supplementary File 1), illustrating that the single passage in mice did not generate further significant changes. There were 13 different amino acid changes and two deletions in the spike protein across the five MA viruses (Fig. 2a, b). All these changes have previously been identified in human isolates (Fig. 2c). The changes are described in detail, with structural information in Supplementary Figs. 1-3. The Q498H and Q493R substitutions (Fig. 2a) have been identified previously for mouse adapted viruses and increase affinity for mACE2^12-17^. The latter contention is supported by our modeling (Supplementary Fig. 2). The non-spike changes comprise nine different amino acid substitutions in Orf1ab (Fig. 2a); with all these changes also identified in human isolates (Supplementary Fig. 4).

**Figure 2.**
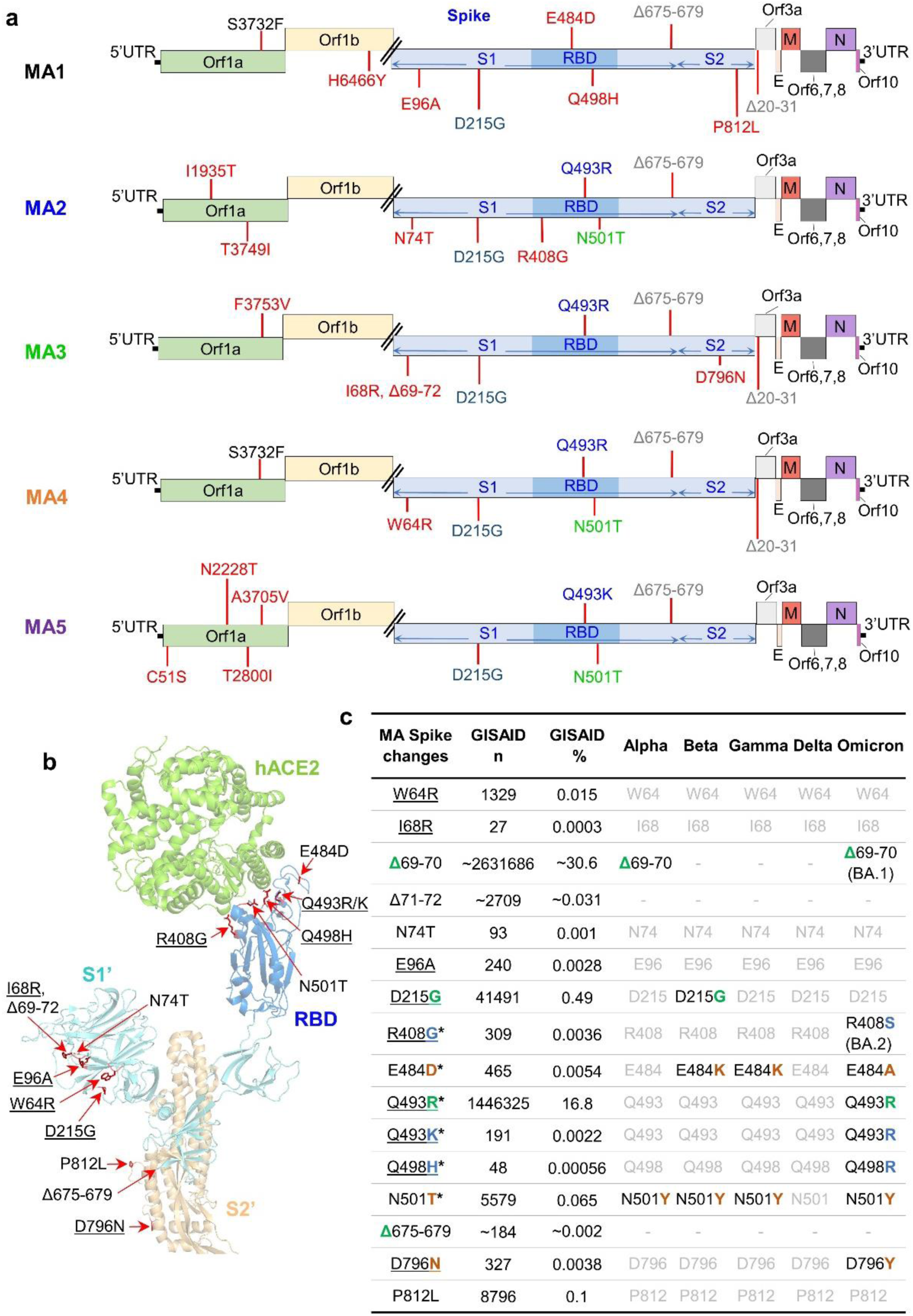
Sequencing of MA viruses. **a** Amino acid changes and deletions (Δ) in MA1, MA2, MA3, MA4 and MA5 (full dataset in Supplementary File 1). Amino acid changes that only appear in one MA virus are in red, with other colors used to show amino acid changes common between at least two MA viruses. **b** Amino acids changes and deletions in any of the MA viruses shown on the structure of SARS-CoV-2_QLD02_ spike bound to hACE2 (PDB: 7DF4). Underlining indicates non-conservative amino changes. **c** Spike changes in the MA viruses. * - amino acid changes located within the RBD. Underlining indicates non-conservative amino acid changes. GISAID n – number of GISAID submissions that contain this change; GISAID % - percentage of all GISAID submissions with this change. Alpha, Beta, Gamma, Delta, Omicron: black text - hallmark changes in variants of concern; grey text –not a hallmark change in variants of concern. Match of amino acid changes in MA viruses with hallmark changes in variants of concern (green - exact match, blue - conservative, brown - non-conservative change).

### MA1 replicates efficiently in several ACE2-negative cell lines

*In vitro* growth kinetics of MA1 and MA2 were studied in more detail using stock virus derived from infected mouse lungs and prepared in HEK293T-mACE2 cells (Fig. 1a). As expected, MA1 and MA2, as well as SARS-CoV-2_QLD02,_ and Alpha and Beta strain viruses, all replicated in HEK293T-hACE2 cells (Fig. 3a). MA1, MA2 and Alpha, Beta, and Omicron strain viruses, but not SARS-CoV-2_QLD02_, replicated in HEK293T-mACE2 cells (Fig. 3b); consistent with Fig. 1b-d and the reported ability of Alpha, Beta and Omicron variants to replicate in mice^6^. Perhaps of interest, given the postulated rodent origins of Omicron^24,25^, our Omicron isolate replicated more efficiently in HEK293T-mACE2 than in HEK293T-hACE2 cells (Fig. 3a, b).

**Figure 3.**
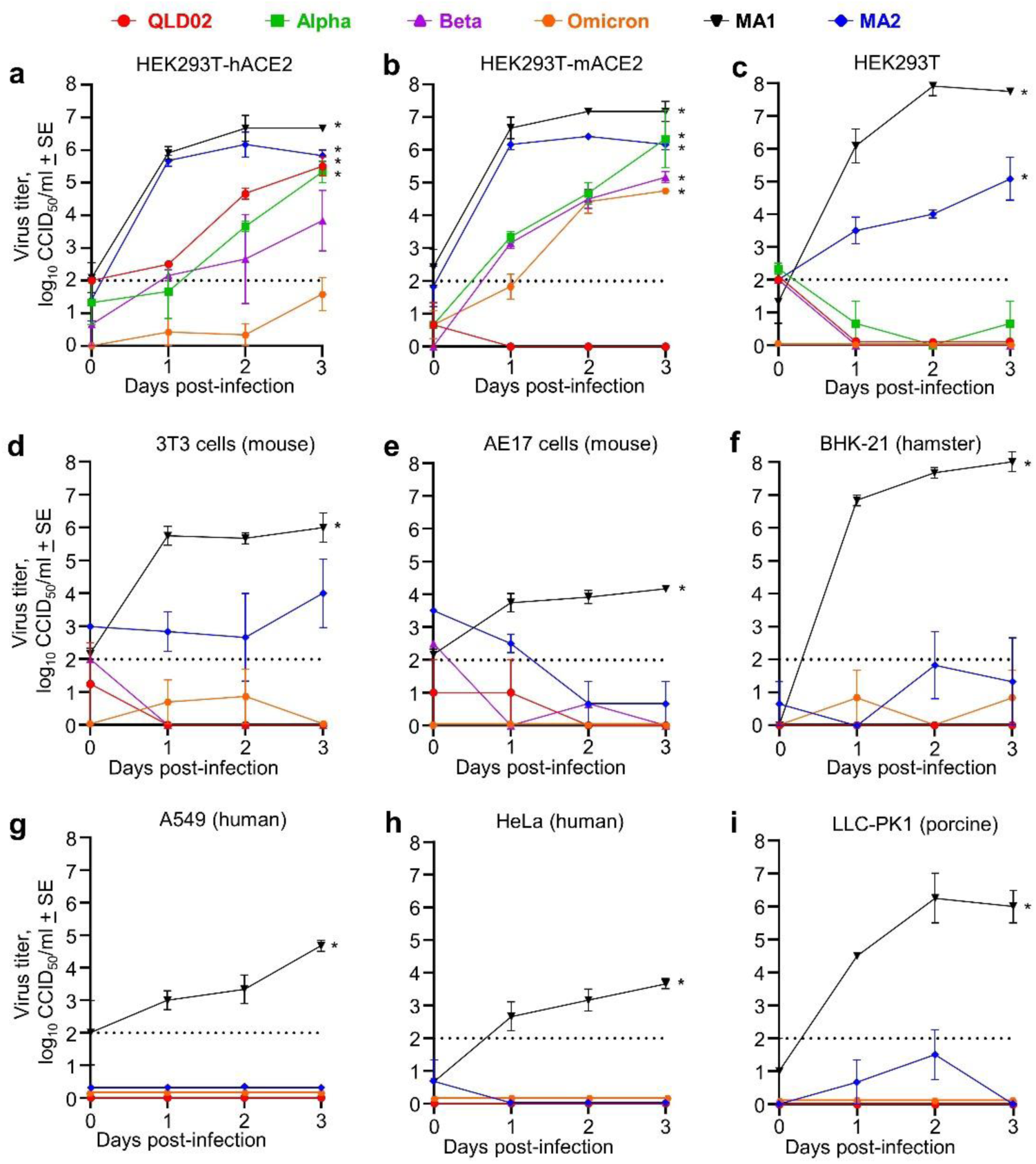
*In vitro* growth kinetics of mACE2-adapted viruses reveal an ACE2-independent entry mechanism. Growth kinetics of the indicated viruses in HEK293T-hACE2 cells **(a)**, HEK293T-mACE2 cells **(b)**, untransduced HEK293T cells **(c)**, 3T3 cells **(d)**, AE17 cells **(e)**, BHK-21 cells **(f)**, A549 cells **(g)**, HeLa cells **(h)** and LLC-PK1 **(i)** cells after infection at MOI = 0.1. n = 3-6 replicates per virus strain per cell line. Dotted line – limit of detection. * p<0.05; statistics by Kolmogorov–Smirnov test for 3 dpi versus 0 dpi.

Remarkably, when replication was examined in control untransduced HEK293T cells, both MA1 and MA2 showed significant replication, with MA1 replicating to ≈8 log_10_CCID_50_/ml in 2 days (Fig. 3c). None of the other viruses were able to replicate efficiently in untransduced HEK293T cells (Fig. 3c), consistent with previous reports^2,27,28^. Replication was also tested in a series of cell lines (Fig. 3d-i) in which (i) publically available RNA-Seq data showed negligible levels of ACE2 mRNA expression (Supplementary Fig. 5a) or (ii) surface expression of ACE2 is absent (LLC-PK1)^29^. MA1 was able to replicate in all the aforementioned cell lines (Fig. 3d-i). In contrast, MA2 and SARS-CoV-2_QLD02_ were unable to replicate in any of these cell lines (Fig. 3d-i); the SARS-CoV-2_QLD02_ results are consistent with previous reports^2,29-32^. Overall, the data suggests ACE2-independent infection by MA1 is robust across multiple cell lines and is functional, at least *in vitro*, across 4 different species (human, mouse, hamster and pig).

None of these cell lines expressed TMPRSS2 mRNA (Supplementary Fig. 5b), with virus fusion instead likely to rely on cathepsin L^33^, with cathepsin L mRNA abundantly expressed in all the cell lines (except BHK-21 which does not have an annotated orthologue) (Supplementary Fig. 5c).

### MA1 and MA2 infection of C57BL/6J mice

To characterize the behavior of MA1 and MA2 during mouse infections, C57BL/6J mice were infected by intrapulmonary inoculation (via the intranasal route) with 10^5^ CCID_50_ MA1 or MA2 per mouse. Virus titers in lungs reached 6-8 log_10_ CCID_50_/g on day 2, with no significant differences seen between MA1 and MA2 (Fig. 4a). These results are broadly similar to those reported previously for mouse adapted viruses^12,17^. Nasal turbinate titers on day 2 were significantly higher for MA1 (≈7 log_10_ CCID_50_/g), when compared with MA2 (Fig. 4b). MA1 and MA2 virus titers were below the level of detection for all other mice and tissues tested (similar to other reports^14^), with the exception of heart where MA1 infection resulted in detectable heart infection on day 2 in 2/4 mice (Fig. 4c). (Infectious virus can also be detected in the heart of K18-hACE mice – Supplementary Fig. 6a). There was no significant weight loss observed for MA1 or MA2 infected mice (Supplementary Fig. 6b).

**Figure 4.**
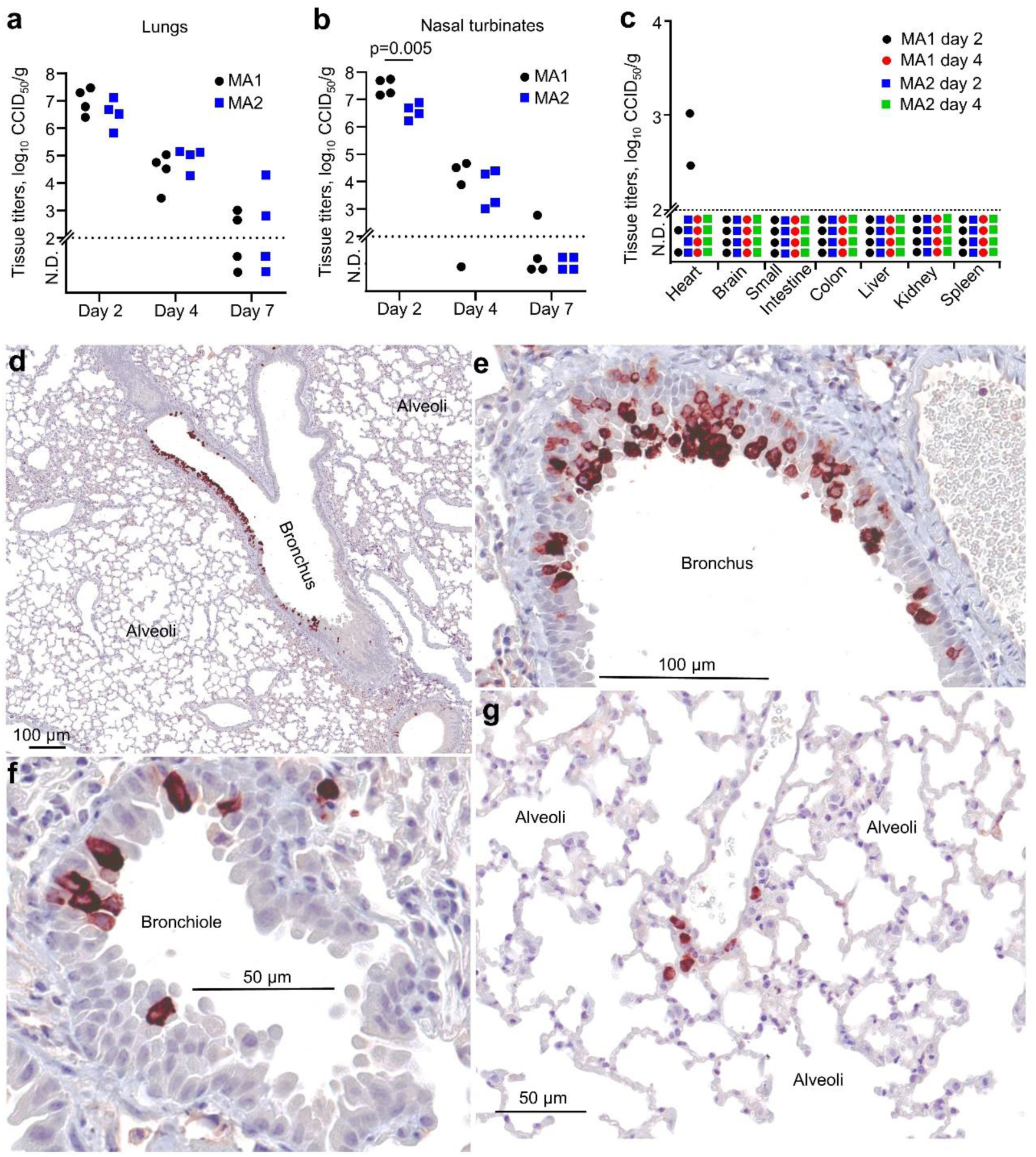
MA1 and MA2 infection in C57BL/6J mice. **a** C57BL/6J mice were infected with MA1 or MA2 and lungs collected at day 2, 4 or 7 post-infection and tissue titers determined by CCID_50_ assays. **b** As for (**a)** for nasal turbinates. Statistics by t-test. **c** As for (**a)** for the indicated tissues. **d-g** IHC using an anti-SARS-CoV-2 spike monoclonal antibody and lungs taken on day 2 after infection of C57BL/6J mice with MA1. Images are representative of lung sections from 4 mice. Dark brown staining (Nova Red) indicates infected cells, often with the expected, clearly discernable, cytoplasmic staining pattern. The large unstained areas in (**d)** and (**g**) are included to illustrate the specificity of the IHC staining.

Immunohistochemistry (IHC) staining with anti-SARS-CoV-2 spike monoclonal antibody showed that MA1 virus infection was localized primarily to the epithelium of bronchi (Fig. 4d-e) and bronchioles (Fig. 4f) in C57BL/6J mouse lungs, with occasional staining of cells in the alveoli (Fig. 4g). A similar picture emerged for MA2 (Supplementary Fig. 7). There was also occasionally prodigious staining of bronchial epithelium in MA1 infected lungs (Supplementary Fig. 8a). Other studies using mouse-adapted SARS-CoV-2 viruses also showed viral staining in bronchial epithelium^14,17,34^, and such staining is also seen after infection with non-mouse adapted viruses of mice expressing hACE2 from the mACE2 promoter^35^. Staining of alveoli was not a prominent feature in MA1 (Fig. 4d, g) or MA2 infected (Supplementary Fig. 7) C57BL/6J mice, with similar results reported for SARS-CoV-2 infection of mice expressing hACE2 from the mACE2 promoter^35^. This contrasts markedly with SARS-CoV-2 infection of K18-hACE2 mice, which show widespread infection of alveoli^35^. Occasional staining of MA1 was also observed in tracheal columnar epithelial cells (Supplementary Fig. 8b), indicating upper respiratory tract infection. Omicron viruses similarly show a propensity to replicate in the upper respiratory tract, rather than in lung parenchyma^36-38^.

Mouse-to-mouse (C57BL/6J to K18-hACE2) transmission was not seen for either MA1 or MA2 (Supplementary Fig. 9a,b). Such transmission has been reported for K18-hACE2 to K18-hACE2 mice^39^ and for deer mice^23^. The complete loss of an intact furin cleavage site in MA1 and MA2 may, at least in part, be responsible (Supplementary Fig. 9c), with this site required for transmission in ferrets^40^, as well as pathogenicity in humans^41^. Loss of this site commonly arises after *in vitro* passage of SARS-CoV-2^42^ and provides a built-in safety feature for our MA viruses.

### Lung pathology after MA1 and MA2 infections of C57BL/6J mice

C57BL/6J mice were infected with MA1 and MA2 and lungs harvested on days 0, 2, 4 and 7 post infection and analyzed by hematoxylin and eosin (H&E) staining. Examples of low magnification images of whole lung sections illustrate the loss of alveolar airspace (Fig. 5a), which reached significance by image analysis on day 7 for MA1 infected mice (Fig. 5b). The ratio of blue (nuclear) to red (cytoplasmic) pixels in H&E stained sections is a measure of cellular infiltration^43^, with lungs from MA1 or MA2 infected mice showing significantly higher cellular infiltrates compared to uninfected mice if all days post infection are taken together (Fig. 5c).

**Figure 5.**
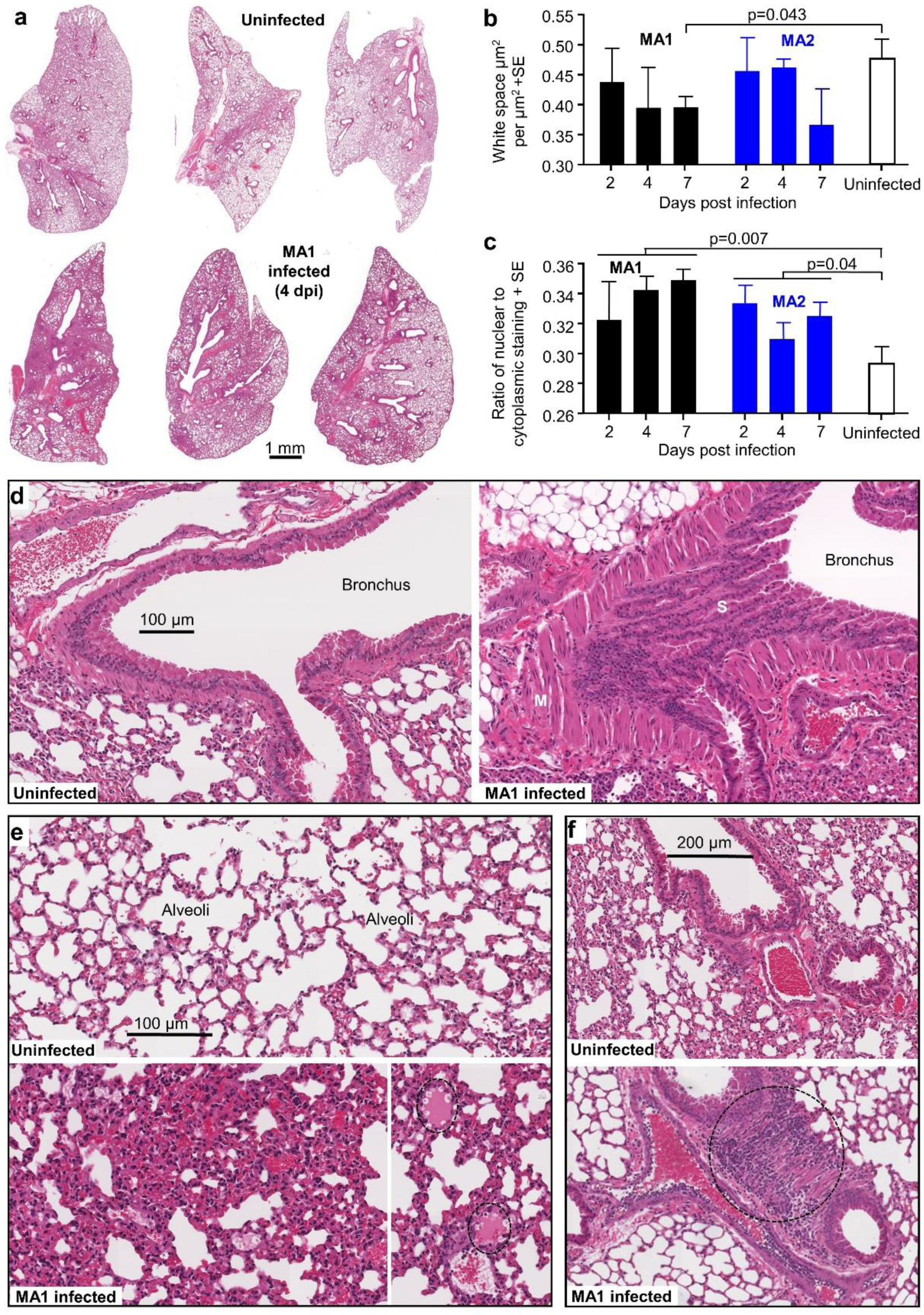
H&E staining of lungs after infection of C57BL/6J mice with MA1 and MA2. **a** Representative low magnification images of H&E stained lung sections for uninfected and MA1 infected C57BL/6J mice taken 4 days post infection. **b** Image analysis to measure lung consolidation quantitated as area of white space (unstained air spaces within the lung) per µm^2^ (n=4 mice per group, with 3 sections scanned per lung and values averaged to produce one value for each lung). Statistics by t-test. **c** Image analysis to quantitate leukocyte infiltration. The ratios of nuclear (blue/dark purple) to cytoplasmic (red) staining of H&E stained lung n=4 mice per group, with 3 sections scanned per lung and values averaged to produce one value for each lung. Statistics by t-tests, with data from the 3 time points for infected mice combined. **d** High magnification image showing bronchus from uninfected mice (left) and MA1 infected mice (right) on day 4 post infection. The latter shows sloughing of the bronchial epithelium (S) and smooth muscle hyperplasia (M). **e** High magnification image showing uninfected lungs (top) and MA1 infected lungs (4 dpi) (bottom). The latter shows lung consolidation and loss of alveolar air spaces, as well as pulmonary edema (dotted ovals, bottom right). **f** High magnification images of uninfected lungs (top) and in MA1-infected lungs on 4 dpi (bottom). The latter shows dense cellular infiltrate (dotted circle).

H&E staining revealed clear evidence of bronchiolar sloughing and smooth muscle hyperplasia in MA1-infected mice (Fig. 5d). The collapse of alveolar spaces, with some pulmonary edema is also evident (Fig. 5e) in MA1 infected lungs, consistent with Fig. 5b. Areas with dense cellular infiltrates were occasionally pronounced in MA1-infected lungs (Fig. 5e, dotted circle), consistent with Fig. 5c. Overall similar lung pathology was seen in MA2-infected mice (Supplementary Fig. 10). MA infection of C57BL/6J mice results in robust lung pathology, with no overtly unique features when compared, for instance, with the infection of K18-hACE2 mice with SARS-CoV-2_QLD02_^9^.

### Inflammatory responses in MA1 infected C57BL/6J mouse lungs are similar to, but less severe than, SARS-CoV-2_QLD02_ infection of K18-hACE2 mouse lungs

RNA-Seq was used to determine the inflammatory responses in C57BL/6J mouse lungs on day 4 post MA1 infection (full gene list in Supplementary File 2a). Differentially expressed genes (DEGs) (n = 1027, Supplementary File 2b) were analyzed for ‘UpStream Regulators’ (USRs) (Supplementary File 2c) and ‘Diseases and Functions’ (Supplementary File 2d) tools of the Ingenuity Pathway Analysis (IPA) software. The same analyses were performed for day 4 lungs from K18-hACE2 mice infected with SARS-CoV-2_QLD02_^44^ (n=1349 DEGs; Supplementary File 2e-h, respectively). A highly significant correlation emerged for cytokine USRs (Fig. 6a; Supplementary File 2i). A number of USR annotations (Fig. 6a, red) indicated that inflammation in K18-hACE2 is more severe (i) with higher z scores for the pro-inflammatory cytokine TNF, and osteopontin (SPP1), a proposed severity marker of COVID-19 ^45^, and (ii) lower z-scores for anti-inflammatory cytokines IL10 and IL37, and Secretoglobin family 1a member 1 (SCGB1A1), a pulmonary surfactant protein that blunts alveolar macrophage responses and mitigates against cytokine surges^46^.

**Figure 6.**
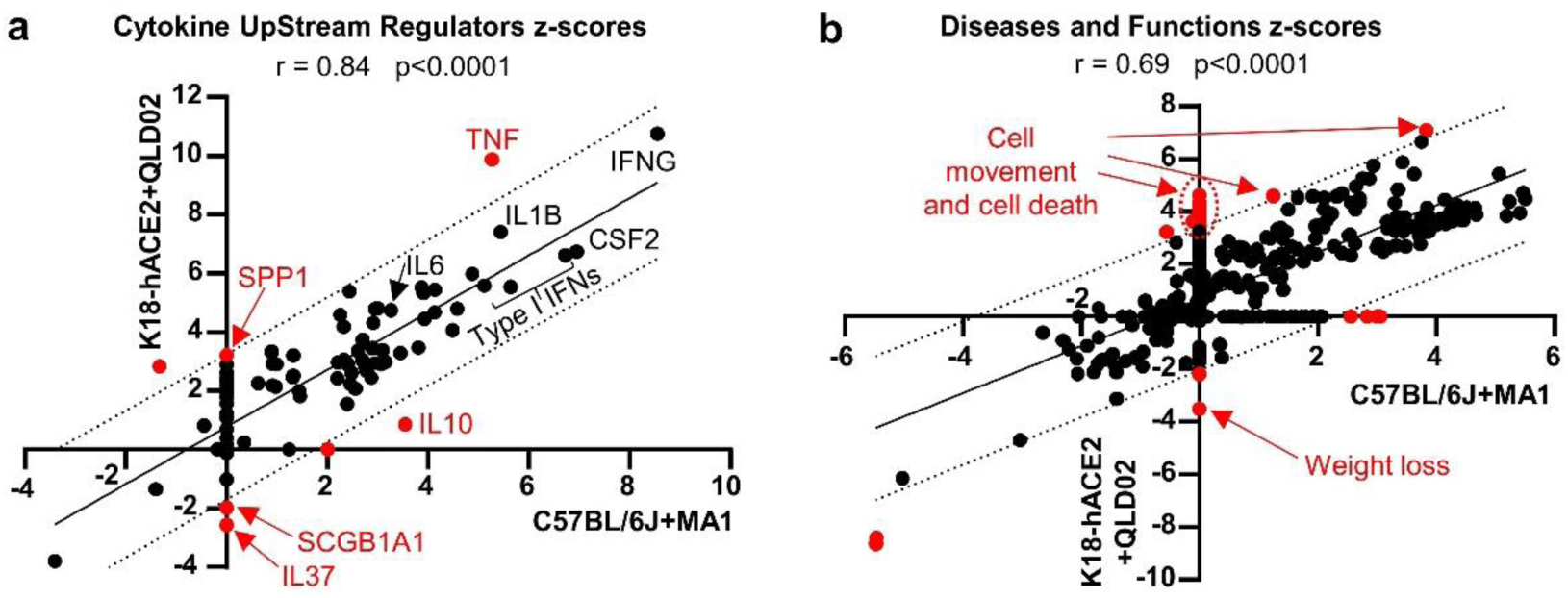
RNA-Seq comparison of MA1 infected C57BL/6J versus SARS-CoV-2_QLD02_ infected K18-hACE2 mouse lungs. RNA-Seq was performed on mouse lungs 4 dpi for MA1 infection of C57BL/6J mice, and SARS-CoV-2_QLD02_ infection of K18-hACE2 mice. DEGs (n = 1027 for C57BL/6J, n = 1349 for K18-hACE2) were analyzed by Ingenuity Pathway Analysis, and z-scores for significant UpStream Regulators **(a)** or Diseases and Functions **(b)** were plotted with C57BL/6J+MA1 on x-axis and K18-hACE2+QLD02 on the y-axis. Statistics by Pearson correlations. Dotted black lines - 95% confidence limits. Solid black line - line of best fit. Annotations outside the 95% confidence limits are colored in red. Selected annotations are labelled (red text). Full data sets are provided in Supplemental File 2.

The Disease and Functions annotation z scores also showed a highly significant correlation, although K18-hACE2 annotations again indicated a more severe inflammatory response, with increased cell movement, cell death, and weight loss (Fig. 6b; Supplementary File 2j), consistent with histology^9^ (Fig. 5) and body weight measurements^44^ (Supplementary Fig. 6b). Thus although MA1 can infect cells lacking ACE2, the inflammatory responses in C57BL/6J mice were very similar to those seen in the K18-hACE2/SARS-CoV-2_QLD02_ model, which is dependent on hACE2.

## DISCUSSION

Herein we describe the rapid *in vitro* adaptation of an original isolate of SARS-CoV-2 to produce viruses capable of infecting mice, with one virus (MA1) also capable of ACE2-independent infection. The latter was robust across multiple cell lines and was functional in at least 4 different species (human, mouse hamster and pig). Our study again highlights the ability of SARS-CoV-2 to evolve rapidly in the face of selection pressure^47^, a feature that has also been documented in individual patients^48^. Our study again highlights the relatively small number of changes (that include Q498H, Q493R/K, and N501T; Supplementary Fig. 1) that allow SARS-CoV-2 to jump species^2,12-14,16,17^.

The ability to infect cells in an ACE2-independent manner, would appear to require the E484D substitution, identified herein and previously^26^. This substitution has been identified in multiple human isolates, in all variants of concern and in a large range of geographical locations around the world (Supplementary Fig. 11). Omicron has an E484A substitution in this position, with Omicron requiring ACE2 expression for infection (Fig. 3)^36^. Modeling suggests the E484D substitution retracts the carboxyl group located between Y489 and F490 of the RBD (Supplementary Fig. 12), perhaps thereby providing a novel binding site. A series of alternative or additional receptors have been reported for SARS-CoV-2 Wuhan strain and include DC-SIGN/L-SIGN^49^, Kidney Injury Molecule-1/T cell immunoglobulin mucin domain 1 (KIM-1/TIM-1)^29^, AXL Receptor Tyrosine Kinase^28^, neuropilin 1 (NRP1)^27^ and CD147^50^. Analysis of publically available RNA-Seq data indicated that only CD147 was expressed at high levels in all cell lines tested herein (Supplementary Fig. 13). However, Wang *et al*. reported replication of an original isolate of SARS-CoV-2 in BHK-21 cells only when human CD147 was over-expressed^50^. The MA viruses are likely to enter via the endosomal route, given the loss of the furin cleavage site^37,51^. This loss improves cleavage by Cathepsin L, which substitutes for TMPRSS2 by cleaving S2’ in endosomes and releasing viral RNA into the cytoplasm^33^. Omicron, for different unknown reasons, shows reduced reliance on TMPRSS2, and relies more on endosomal entry^36,52,53^. The D614G change, which is present in all SARS-CoV-2 variants (except the original Wuhan strain), also increases entry via the cathepsin L route^33^. Cathepsin L is widely expressed in human nasal and lung epithelial cells^33^ and in the cell lines tested herein (Supplementary Fig. 5c).

The concept that Omicron might have arisen via rodents^24,54^ is perhaps supported by the observation that Omicron replicated more efficiently in mACE2-expressing cells when compared with hACE2-expressing cells (Fig. 3a,b). Omicron binding to mACE2 is significantly higher than Alpha and Beta variants, which are also able to use mACE2 as a receptor^55^. Increased mACE2-binding by Omicron may involve the Q493R, Q498R and N501Y substitutions (Fig. 2c, Omicron), given these changes are selected in mouse-adapted viruses^12-15,17^. Our MA viruses have Q493R, Q498H and N501T substitutions (Fig. 2c; Supplementary Fig. 1), with modeling suggesting Q493R and Q498H increase mACE2 binding (Supplementary Fig. 2). Finally, like Omicron^36^, MA1, MA2 and other mouse adapted viruses^14,17^, show a preference for replication in the upper respiratory tract.

The pathological consequences of ACE-2 independent infection remain to be characterized, and one might speculate that ACE-independent infection may play a role in long-COVID^56,57^. Our studies of acute MA1 infections in C57BL/6J mice argue that when ACE2-dependent infection remains possible, the ability also to infect via an ACE2-independent mechanism does not result in overtly unique pathology or immunopathology. Both the development of mouse models of long COVID, evaluation of infection and disease in ACE2^-/-^ mice and the identification of the alternate receptor used by MA1 are thus the focus of our future studies.

## ACKNOWLEDGEMENTS

From QIMR Berghofer Medical Research Institute we thank Dr I Anraku for managing the PC3 (BSL3) facility, animal house staff for mouse breeding and agistment, Paul Collins for library preparation and RNA-Seq, and Dr Clay Winterford, Sang-Hee Park and Crystal Chang for the histology and immunohistochemistry. We thank Dr Alyssa Pyke and Mr Fredrick Moore (Queensland Health, Brisbane) for providing the SARS-CoV-2 isolates. We thank Monash Genome Modification Platform for providing the plasmid containing the mouse-codon optimized human ACE2 gene. We thank Clive Berghofer and the Brazil Family Foundation (and many others) for their generous philanthropic donations to support SARS-CoV-2 research at QIMR Berghofer MRI. The project was also partly funded by an intramural QIMRB seed grant awarded to D.J.R. A.S. holds an Investigator grant from the National Health and Medical Research Council (NHMRC) of Australia (APP1173880).

## AUTHOR CONTRIBUTIONS

Conceptualization, D.J.R.; Methodology, D.J.R. and A.S.; Formal analysis, D.J.R., A.S., T.D., C.B.; Investigation, D.J.R., K.Y., T.T.L., and B.T.; Resources, A.S.; Data curation, D.J.R., A.S., and T.D.; Writing – D.J.R., AS; Visualization, D.J.R., A.S., and T.D.; Supervision, D.J.R. and A.S., Project administration, D.J.R. and A.S.; Funding acquisition, A.S. and D.J.R.

## DECLARATION OF INTERESTS

The authors declare no competing interests.

## DATA AVAILABILITY

All raw sequencing data (fastq files) are available from the Sequence Read Archive (SRA), BioProject accession: PRJNA804321. All other data is available within the paper and supporting information files.

## MATERIALS and METHODS

### Ethics statement and regulatory compliance

All mouse work was conducted in accordance with the “Australian code for the care and use of animals for scientific purposes” as defined by the National Health and Medical Research Council of Australia. Mouse work was approved by the QIMR Berghofer Medical Research Institute Animal Ethics Committee (P3600, A2003-607). For intrapulmonary inoculations, mice were anesthetized using isoflurane. Mice were euthanized using CO_2_ or cervical dislocation. Obtaining samples from patients and using human SARS-CoV-2 isolates was approved by the QIMR Berghofer MRI Human Research Ethics Committee (P3600).

Breeding and use of GM mice was approved under a Notifiable Low Risk Dealing (NLRD) Identifier: NLRD_Suhrbier_Oct2020: NLRD 1.1(a). Cloning and use of lentiviral vectors for transduction of ACE2 into cell lines was approved under an NLRD (OGTR identifier: NLRD_Suhrbier_Feb2021: NLRD 2.1(l), NLRD 2.1(m)).

All infectious SARS-CoV-2 work was conducted in a dedicated suite in a biosafety level-3 (PC3) facility at the QIMR Berghofer MRI (Australian Department of Agriculture, Water and the Environment certification Q2326 and Office of the Gene Technology Regulator certification 3445). All work was approved by the QIMR Berghofer Medical Research Institute Safety Committee (P3600).

### Cell lines and SARS-CoV-2 culture

Vero E6 (C1008, ECACC, Wiltshire, England; obtained via Sigma Aldrich, St. Louis, MO, USA), Lenti-X 293T (Takara Bio), AE17 (a gift from Dr Delia Nelson, Faculty of Health Sciences, Curtin Medical School), NIH-3T3 (American Type Culture Collection, ATCC, CRL-1658), LLC-PK1 (a gift from Prof. Roy Hall, University of Queensland), A549 (ATCC CCL-185), BHK-21 (ATCC# CCL-10) and HeLa (ATCC-CLL 2) cells were cultured in medium comprising DMEM for Lenti-X 293T and A549 cells, M199 for LLC-PK1 cells or RPMI1640 for all the others (Gibco), supplemented with (endotoxin free^58^) 10% fetal bovine serum (FBS), penicillin (100 IU/ml)/streptomycin (100 μg/ml) (Gibco/Life Technologies) and L-glutamine (2 mM) (Life Technologies). Cells were cultured at 37°C and 5% CO_2_. Cells were routinely checked for mycoplasma (MycoAlert Mycoplasma Detection Kit MycoAlert, Lonza) and FBS was assayed for endotoxin contamination before purchase^58^.

The SARS-CoV-2 original, Alpha and Beta isolates were kindly provided by Dr Alyssa Pyke (Queensland Health Forensic & Scientific Services, Queensland Department of Health, Brisbane, Australia). The viruses (hCoV-19/Australia/QLD02/2020, Alpha variant hCoV-19/Australia/QLD1517/2021 and Beta variant hCoV-19/Australia/QLD1520/2020) were isolated from patients and sequences deposited at GISAID (https://www.gisaid.org/). Virus stocks were generated by infection of Vero E6 cells at multiplicity of infection (MOI)≈0.01, with supernatant collected after 2-3 days, cell debris removed by centrifugation at 3000 x g for 15 min at 4°C, and virus aliquoted and stored at -80°C. Virus titers were determined using standard CCID_50_ assays (see below). The virus was determined to be mycoplasma free using co-culture with a non-permissive cell line (i.e. HeLa) and Hoechst staining as described^59^.

An Omicron isolate was isolated from a SARS-CoV-2 infected individual in Brisbane Australia. A nasal swab was acquired, submerged and agitated in 1 ml RPMI + 10% FBS + penicillin (100 IU/ml)/streptomycin (100 μg/ml) (Gibco/Life Technologies), and was transported to the laboratory on ice. After centrifugation at 20,000 x g for 10 min the supernatant was cultured on 2×10^6^ Vero E6-TMPRSS2 cells^9^ in a T25 flask for 3 days at 37°C and 5% CO_2_; 1 ml was passaged on to a confluent T75 flask of Vero E6-TMPRSS2 cells, and supernatant harvested and aliquoted after 3 days. Virus titers were determined using standard CCID_50_ assays (see below). For sequencing, viral RNA was isolated from infected Vero E6-TMPRSS2 cells by TRIzol extraction as per manufacturer’s instructions. Reverse transcription was performed using ProtoScript® II First Strand cDNA Synthesis Kit (New England Biolabs) as per manufacturer’s instructions. PCR of a spike gene fragment was performed using Q5 High-Fidelity 2X Master Mix (New England Biolabs) and the following primers: Forward 5’ – TTGAACTTCTACATGCACCAGC – 3’ and Reverse 5’ – CCAGAAGTGATTGTACCCGC – 3’. The fragment was gel purified using Monarch DNA Gel Extraction Kit (New England Biolabs), and Sanger sequencing was performed using BigDye Terminator v3.1 and either the forward or the reverse primer. The virus was confirmed as the Omicron variant due to containing the following amino acid changes compared to SARS-CoV-2_QLD02_; N547K, D614G, H655Y, N679K, P681H, N764K, D796Y, N856K.

### CCID_50_ assays

Vero E6 cells were plated into 96 well flat bottom plates at 2×10^4^ cells per well in 100 µl of medium. For tissue titer, tissue was homogenized in tubes each containing 4 ceramic beads twice at 6000 x g for 15 seconds, followed by centrifugation twice at 21000 x g for 5 min before 5 fold serial dilutions in 100 µl RPMI1640 supplemented with 2% FBS. For cell culture supernatant, 10 fold serial dilutions were performed in 100 µl RPMI1640 supplemented with 2% FBS. 100 µl of serially diluted samples were added to Vero E6 cells and the plates cultured for 5 days at 37°C and 5% CO_2_. The virus titer was determined by the method of Spearman and Karber.

### SARS-CoV-2 passaging in hACE2 and mACE2 co-cultures

Lentivirus encoding hACE2, mACE2 or mACE2^N31K/H353K^ were produced in HEK293T cells by plasmid transfection and were used to transduce HEK293T cells, as described previously^2^. HEK293T-hACE2 or HEK293T-mACE2^N31K/H353K^ were mixed with equal cell numbers of HEK293T-mACE2, and 500,000 cells (i.e. 250,000 HEK293T-hACE2 or mACE2^N31K/H353K^ + 250,000 HEK293T-mACE2) were seeded in 6 well plates overnight. Co-cultures were infected with SARS-CoV-2 at MOI≈0.1. Every 3-4 days, supernatant was collected and centrifuged at 2000 x g at 4°C for 5 min, and 200 µl was added to fresh co-cultures and the remaining was stored at -80°C. This was performed a total of 9 times before 1 ml was passaged on to HEK293T-mACE2 cells. After one passage in HEK293T-mACE2 cells, supernatant was used to infect new HEK293T-mACE2 cells for 2 hrs, then inoculum was removed and cells were washed 3 times with PBS and media replaced. Supernatant was harvested at 0 hr and 72 hrs post infection for virus titration by CCID_50_ to confirm virus replication in mACE2 expressing cells. Virus was then passaged one additional time in HEK293T-mACE2 cells and supernatant stored at -80°C for use in sequencing and mouse infections.

For growth kinetics experiments, HEK293T, HEK293T-hACE2 and HEK293T-mACE2, NIH-3T3, AE17, BHK-21, A549, HeLa or LLC-PK1 cells were infected with SARS-CoV-2 (QLD02, MA1, MA2, Alpha, Beta or Omicron) at MOI≈0.1 for 1 hr at 37°C, cells were washed with PBS and media replaced. Culture supernatant was harvested at the indicated time points and titered by CCID_50_ assay as described above.

### Mouse intrapulmonary SARS-CoV-2 infection

Female C57BL/6J mice (∼ 6 months old at the time of infection) were purchased from Animal Resources Centre (Canning Vale, WA, Australia). The conditions the mice were kept are as follows: light = 12:12 h dark/light cycle, 7:45 a.m. sunrise and 7:45 p.m. sunset, 15 min light dark and dark light ramping time. Enclosures: M.I.C.E cage (Animal Care Systems, Colorado, USA). Ventilation: 100% fresh air, eight complete air exchange/h/rooms. In-house enrichment: paper cups (Impact-Australia); tissue paper, cardboard rolls. Bedding: PuraChips (Able scientific) (aspen fine). Food: Double bagged norco rat and mouse pellet (AIRR, Darra, QLD). Water: deionized water acidified with HCl (pH = 3.2). Mice were anesthetized using isoflurane and given an intrapulmonary inoculation ≈5×10^4^ CCID_50_ SARS-CoV-2 delivered via the intranasal route in 50 µl. Mice were sacrificed by cervical dislocation at day 2, 4 or 7 and lungs, nasal turbinates, brain, small intestine, colon, liver, kidney and spleen were collected. Right lung and all other organs were immediately homogenized in tubes each containing 4 beads twice at 6000 x g for 15 seconds, and used in tissue titration as described above. Left lungs were fixed in 10% formalin for histology.

K18-hACE2 mice (strain B6.Cg-Tg(K18-ACE2)2Prlmn/J, JAX Stock No: 034860)^60^ were purchased from The Jackson Laboratory, USA, and bred and maintained in-house at QIMRB as heterozygotes by crossing with C57BL/6J mice^10^. Mice were genotyped using Extract-N-Amp Tissue PCR Kit (Sigma Aldrich) according to manufacturers’ instructions with the following primers; Forward 5’-CTTGGTGATATGTGGGGTAGA-3’ and Reverse 5’-CGCTTCATCTCCCACCACTT-3’ (recommended by NIOBIOHN, Osaka, Japan). Thermocycling conditions were as follows; 94°C 3 min, 35 cycles of 94°C 30 s, 55.8 °C 30 s, 72°C 1 min, and final extension of 72°C 10 min.

### RNA sequencing

For viral RNA purification, cell culture supernatants were processed using the NucleoSpin RNA Virus kit (Machery-Nagel) as per manufacturers’ instructions. For mouse lung RNA isolation, lungs were transferred from RNAlater to TRIzol (Life Technologies) and were homogenized twice at 6000 x g for 15 sec. Homogenates were centrifuged at 14,000 × g for 10 min and RNA was isolated as per manufacturers’ instructions.

RNA concentration and quality was measured using TapeStation D1K TapeScreen assay (Agilent). cDNA libraries were prepared using the Illumina TruSeq Stranded mRNA library prep kit and the sequencing performed on the Illumina Nextseq 550 platform generating 75 bp paired end reads.

For virus sequences, per base sequence quality for >90% bases was above Q30 for all samples. The quality of raw sequencing reads was assessed using FastQC^61^ (v0.11.80), and trimmed using Cutadapt^62^ (v2.3) to remove adapter sequences and low-quality bases. Trimmed reads were aligned using STAR^63^ (v2.7.1a) to a SARS-CoV-2 isolate Wuhan-Hu-1 (NC_045512.2; 29903 bp). Aligned reads were viewed using Integrative Genome Viewer (IGV)^64^, and any position with >20% change compared to the reference genome was manually curated. SAMtools mpileup was used to produce a consensus sequence from mapped reads^65^.

For mouse lungs, per base sequence quality for >90% bases was above Q30 for all samples. The quality of raw sequencing reads was assessed using FastQC^61^ (v0.11.80), and trimmed using Cutadapt^62^ (v2.3) to remove adapter sequences and low-quality bases. Trimmed reads were aligned using STAR^63^ (v2.7.1a) to a combined reference that included the mouse GRCm39 primary assembly and the GENCODE M27 gene model^66^, SARS-CoV-2 isolate Wuhan-Hu-1 (NC_045512.2; 29903 bp). Mouse gene expression was estimated using RSEM^67^ (v1.3.0). Reads aligned to SARS-CoV-2 were counted using SAMtools^65^ (v1.9). Differential gene expression in the mouse was analyzed using EdgeR (3.22.3) and modelled using the quasi-likelihood F-test, glmQLFTest().

### Pathway analysis

Up-Stream Regulators (USRs) and Diseases and Functions enriched in differentially expressed genes in direct and indirect interactions were investigated using Ingenuity Pathway Analysis (IPA) (QIAGEN).

### SARS-CoV-2 amino acid change analyses and modelling

The GISAID (https://www.gisaid.org/phylodynamics/global/nextstrain/)^68^ EpiCoV ‘search’ function (https://www.epicov.org/epi3/frontend#577a0f) was used to identify the number of GISAID SARS-CoV-2 sequence submissions containing each amino acid change in the MA viruses. Data was filtered for submissions that selected “Human” as the host. Data was accessed on 21^st^ February 2021 when the number of total submissions was 8,601,773, and this was used to calculate the proportion of each amino acid change in the MA viruses among total GISAID sequence submissions.

PyMOL v4.60 (Schrodinger) was used for mutagenesis of the crystal structure of SARS-CoV-2 spike bound with ACE2 from the protein data bank (7DF4)^69^.

### Lung histopathology and immunohistochemistry

Lungs and trachea (via pluck necropsy. i.e. removal of tongue, larynx, trachea, lungs, heart, and part of the esophagus in one piece) were fixed in 10% formalin, embedded in paraffin, and sections stained with H&E (Sigma Aldrich). Slides were scanned using Aperio AT Turbo (Aperio, Vista, CA USA) and analyzed using Aperio ImageScope software (LeicaBiosystems, Mt Waverley, Australia) (v10) and the Positive Pixel Count v9 algorithm. Automatic quantitation of white space was undertaken using QuPath v0.2.3^70^. Immunohistochemistry for SARS-CoV-2 antigen was undertaken using mouse anti-SARS-CoV-2 spike monoclonal antibody 1E8 (Hobson-Peters *et al*. in preparation) as described previously^2^.

### Statistics

Statistical analyses of experimental data were performed using IBM SPSS Statistics for Windows, Version 19.0 (IBM Corp., Armonk, NY, USA). The t-test was used when the difference in variances was <4, skewness was > -2 and kurtosis was <2. Otherwise, the non-parametric Kolmogorov-Smirnov test was used.

## Notes

### Competing Interest Statement

The authors have declared no competing interest.

