## Supplementary Figures for "ACE2-independent SARS-CoV-2 infection and mouse adaption emerge after passage in cells expressing human and mouse ACE2"

| Spike substitutions and deletions | Present in MA | Notes |
|---|---|---|
| E484D | MA1 | Associated with ACE2-independent infection[1], and changes in this position are implicated in antibody escape[2]. |
| P812L | MA1 | Located near the S2' cleavage site; cleaved by TMPRSS2[3]. |
| Q498H | MA1 | Present in other mouse adapted strains [4-6]. Believed to increase mACE2 affinity (see Supplementary Fig. 2). |
| E96A | MA1 | E96A arose after passage in Vero E6 cells[7]; however, role unknown. |
| R408G | MA2 | Similar substitution in Omicron BA.2 (see Supplementary Fig. 3). Function of this residue is not known. |
| Q493R/Q493K | MA2,3,4,5 | Present in other mouse adapted strains [5,8]. Believed to increase mACE2 affinity (see Supplementary Fig. 2). |
| ∆675-679 | MA1,2,3,4,5 | The deletion of QTQTN removes furin cleavage function and commonly arises after in vitro passage of SARS-CoV-2[9]. RNA-Seq illustrated that no reads showed a functional furin cleavage site for any MA virus (see also Supplementary Fig. 9c). As an intact site is associated with transmission and pathogenicity [3,10,11], this deletion provides a built-in safety feature for these MA viruses. |
| D215G | MA1,2,3,4,5 | This substitution was already present in 11% of the SARS-CoV-$2_{QLD02}$ stock population (Supplementary Fig. 11i). Present in Beta variant. |
| N501T | MA2,4,5 | N501T was selected in all three HEK293T-mACE2 + HEK293T-$mACE2^{N31K/H353K}$ co-cultures, but none of the HEK293T-mACE2 + HEK293T-hACE2 co-cultures. N501T is reported to be selected for in mink and ferrets[12-14]. |
| N74T | MA2 | A substitution that removes a glycosylation site[15]. |
| Δ69-70 | MA3 | Deletion was accompanied by an I68R substitution (Fig. 2a, b), changes similar to the 69-70 deletion that may increase spike incorporation into virions[16]. |
| W64R | MA4 | Previously reported to arise consistently after in vitro passage of spike-pseudotyped vesicular stomatitis virus (VSV) in hACE2-expressing cells[17]. |
| D796N | MA3 | Not described |


1 Puray-Chavez, M. *et al.* Systematic analysis of SARS-CoV-2 infection of an ACE2-negative human airway cell. *Cell Reports* **36**, 109364, (2021).
2 Greaney, A. J. *et al.* Mapping mutations to the SARS-CoV-2 RBD that escape binding by different classes of antibodies. *Nature Communications* **12**, 4196, (2021).
3 Bestle, D. *et al.* TMPRSS2 and furin are both essential for proteolytic activation of SARS-CoV-2 in human airway cells. *Life Science Alliance* **3**, e202000786, (2020).
4 Zhang, Y. *et al.* SARS-CoV-2 Rapidly Adapts in Aged BALB/c Mice and Induces Typical Pneumonia. *Journal of Virology* **95**, e02477-02420, (2021).
5 Huang, K. *et al.* Q493K and Q498H substitutions in Spike promote adaptation of SARS-CoV-2 in mice. *EBioMedicine* **67**, 103381, (2021).
6 Wang, J. *et al.* Mouse-adapted SARS-CoV-2 replicates efficiently in the upper and lower respiratory tract of BALB/c and C57BL/6J mice. *Protein Cell* **11**, 776-782, (2020).
7 Ragan, I. K. *et al.* A Whole Virion Vaccine for COVID-19 Produced via a Novel Inactivation Method and Preliminary Demonstration of Efficacy in an Animal Challenge Model. *Vaccines (Basel)* **9**, 340, (2021).
8 Leist, S. R. *et al.* A Mouse-Adapted SARS-CoV-2 Induces Acute Lung Injury and Mortality in Standard Laboratory Mice. *Cell* **183**, 1070-1085.e1012, (2020).
9 Liu, Z. *et al.* Identification of Common Deletions in the Spike Protein of Severe Acute Respiratory Syndrome Coronavirus 2. *Journal of Virology* **94**, e00790-00720, (2020).
10 Peacock, T. P. *et al.* The furin cleavage site in the SARS-CoV-2 spike protein is required for transmission in ferrets. *Nature Microbiology* **6**, 899-909, (2021).
11 Johnson, B. A. *et al.* Loss of furin cleavage site attenuates SARS-CoV-2 pathogenesis. *Nature* **591**, 293-299, (2021).
12 Welkers, M. R. A., Han, A. X., Reusken, C. B. E. M. & Eggink, D. Possible host-adaptation of SARS-CoV-2 due to improved ACE2 receptor binding in mink. *Virus Evolution* **7**, (2021).
13 Richard, M. *et al.* SARS-CoV-2 is transmitted via contact and via the air between ferrets. *Nature Communications* **11**, 3496, (2020).
14 Zhou, J. *et al.* Mutations that adapt SARS-CoV-2 to mink or ferret do not increase fitness in the human airway. *Cell Reports* **38**, 110344, (2022).
15 Watanabe, Y., Allen, J. D., Wrapp, D., McLellan, J. S. & Crispin, M. Site-specific glycan analysis of the SARS-CoV-2 spike. *Science (New York, N.Y.)* **369**, 330-333, (2020).
16 Meng, B. *et al.* Recurrent emergence of SARS-CoV-2 spike deletion H69/V70 and its role in the Alpha variant B.1.1.7. *Cell Reports* **35**, 109292, (2021).
17 Dieterle, M. E. *et al.* A Replication-Competent Vesicular Stomatitis Virus for Studies of SARS-CoV-2 Spike-Mediated Cell Entry and Its Inhibition. *Cell Host Microbe* **28**, 486-496.e486, (2020).


**Supplementary Fig. 1. Spike amino acid changes in MA viruses.** Detailed description of changes in MA viruses

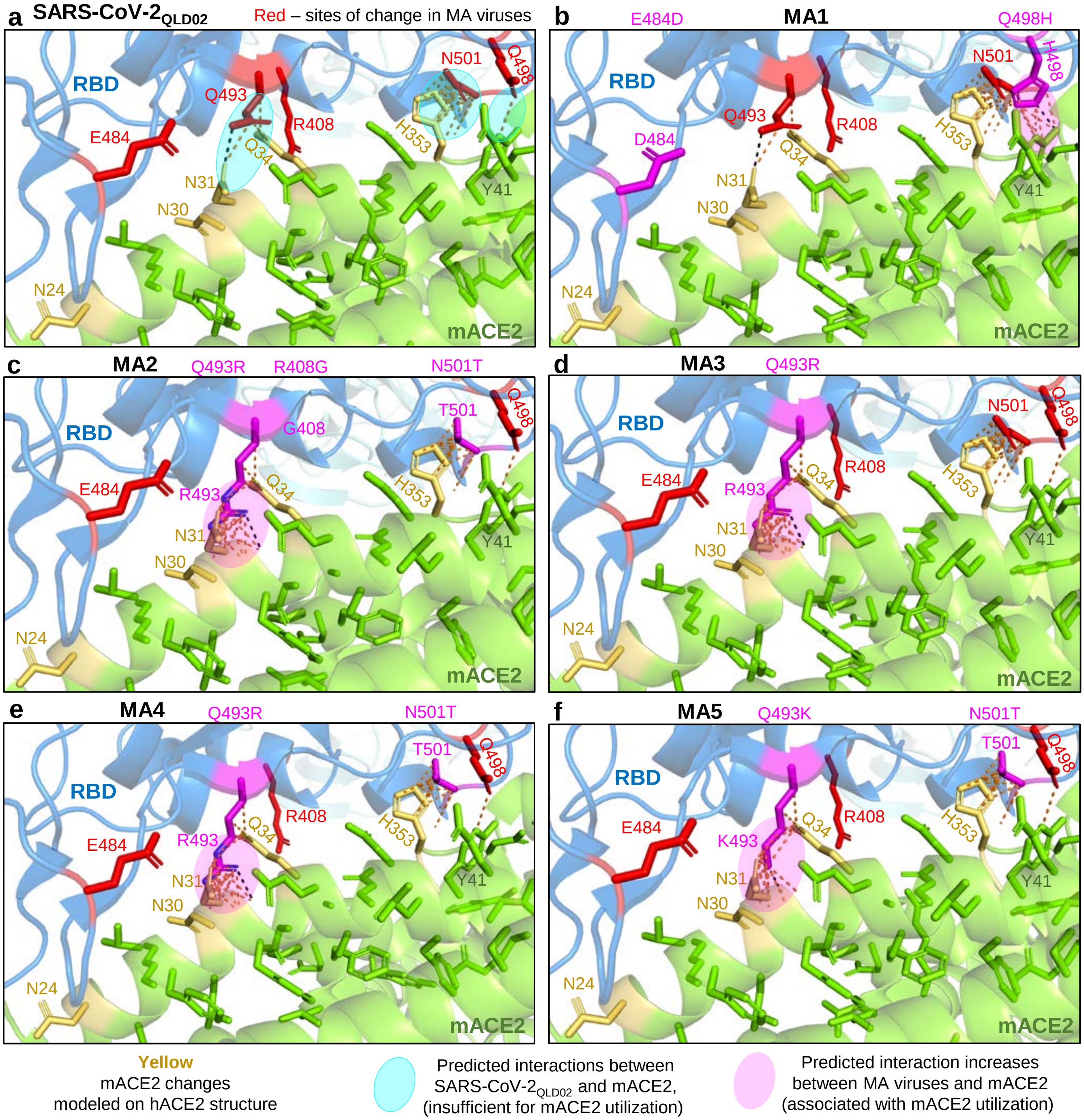


**Supplementary Fig. 2. Modelling RBD changes.** RBD substitutions for MA viruses were modelled using PyMOL to visualize their potential effects on mACE2 binding. **a** Interactions between SARS-CoV-2$_{QLD02}$ RBD and mACE2 are predicted between Q493 and N31/Q34, N501 and H353, and Q498 and Y41 (dashed lines/pale turquoise ovals). These interactions are insufficient to support replication, and are largely retained for MA viruses and mACE2. **b** MA1 has additional predicted interactions between H498 and Y41 (magenta oval). **c-f** MA2-5 have additional predicted interactions between K/R493 and N31/Q34 (**c-f,** magenta ovals).

MA1 and MA2, the viruses with the highest capacity to infect lungs of wild-type mice (Fig. 1d), had two other amino acid changes that modeling predicted are not directly involved in interactions with mACE2, E484D for MA1 and R408G for MA2 (**b, c**) (see also Supplementary Fig. 3).

N501T is a conservative change and modelling suggested that the N501T change would not significantly affect interactions of MA viruses with H353 of mACE2 (**c,e,f**).

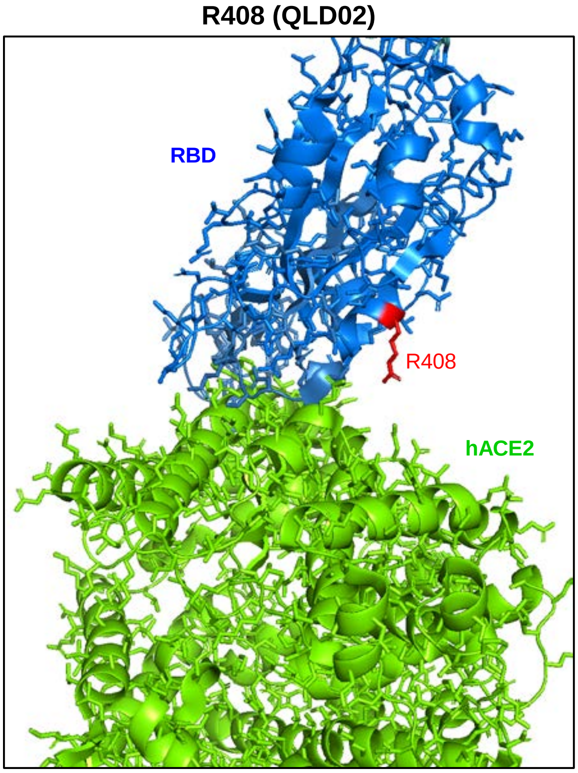


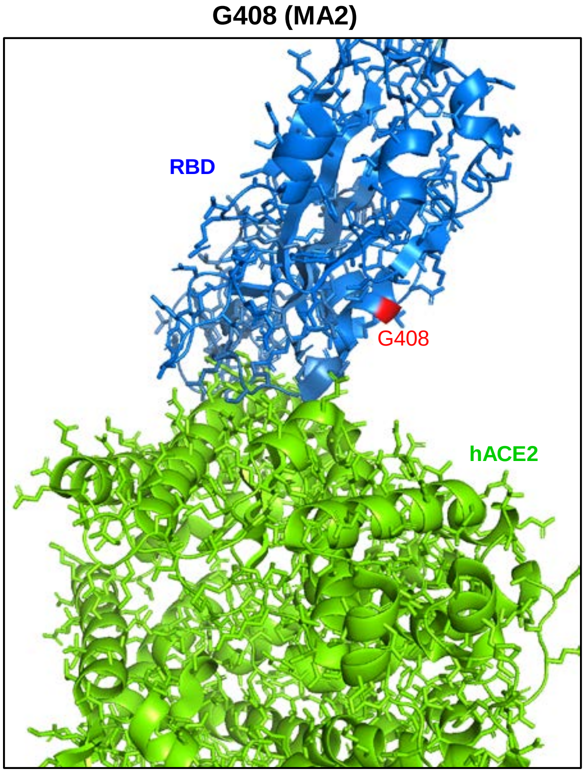


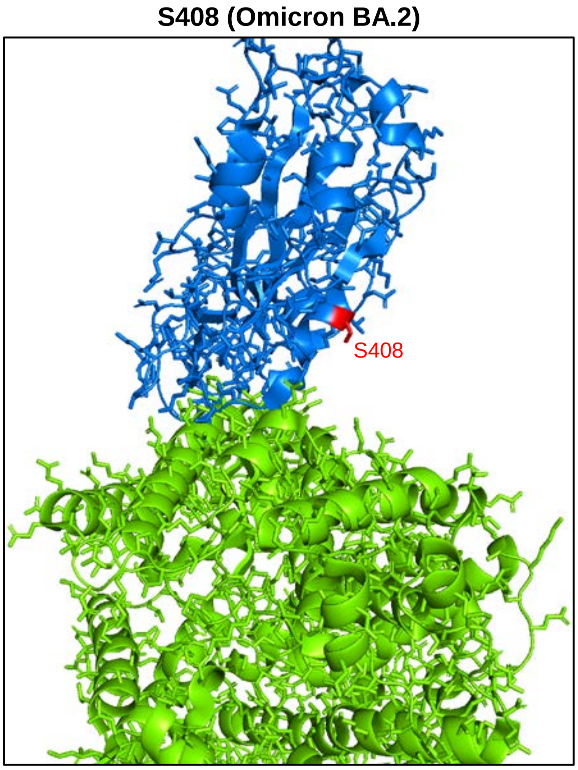


**Supplementary Fig. 3. Position 408 amino acid changes in MA2 and omicron (BA.2).** The R408G change in MA2 and the R408S change in Omicron BA.2 removes a side chain that is predicted to point away from the RBD-hACE2 interface.

The structure of the spike RBD bound to hACE2 (PDB: 7df4) viewed in PyMOL with R408 for QLD02, G408 for MA2, or S408 for Omicron, colored in red. Green = hACE2. Blue = spike RBD.

| Non-spike substitutions; Orf1ab numbering | Non-spike substitutions; Nsp numbering | GISAID isolates n | Percentage of GISAID entries (%) |
|---|---|---|---|
| Orf1ab C51S | nsp1 C51S | 66 | 0.00077 |
| Orf1ab I1935T | nsp3 I1117T | 933 | 0.011 |
| Orf1ab N2228T | nsp3 N1410T | 490 | 0.0057 |
| Orf1ab T2800I | nsp4 T37I | 5292 | 0.062 |
| Orf1ab A3705V | nsp6 A136V | 6546 | 0.076 |
| Orf1ab S3732F | nsp6 S163F | 550 | 0.0064 |
| Orf1ab T3749I | nsp6 T180I | 1700 | 0.02 |
| Orf1ab F3753V | nsp6 F184V | 2525 | 0.03 |
| Orf1ab H6466Y | nsp15 H14Y | 733 | 0.0085 |

**Supplementary Fig. 4. Orf1ab amino acid changes in MA viruses and prevalence in human isolates.**

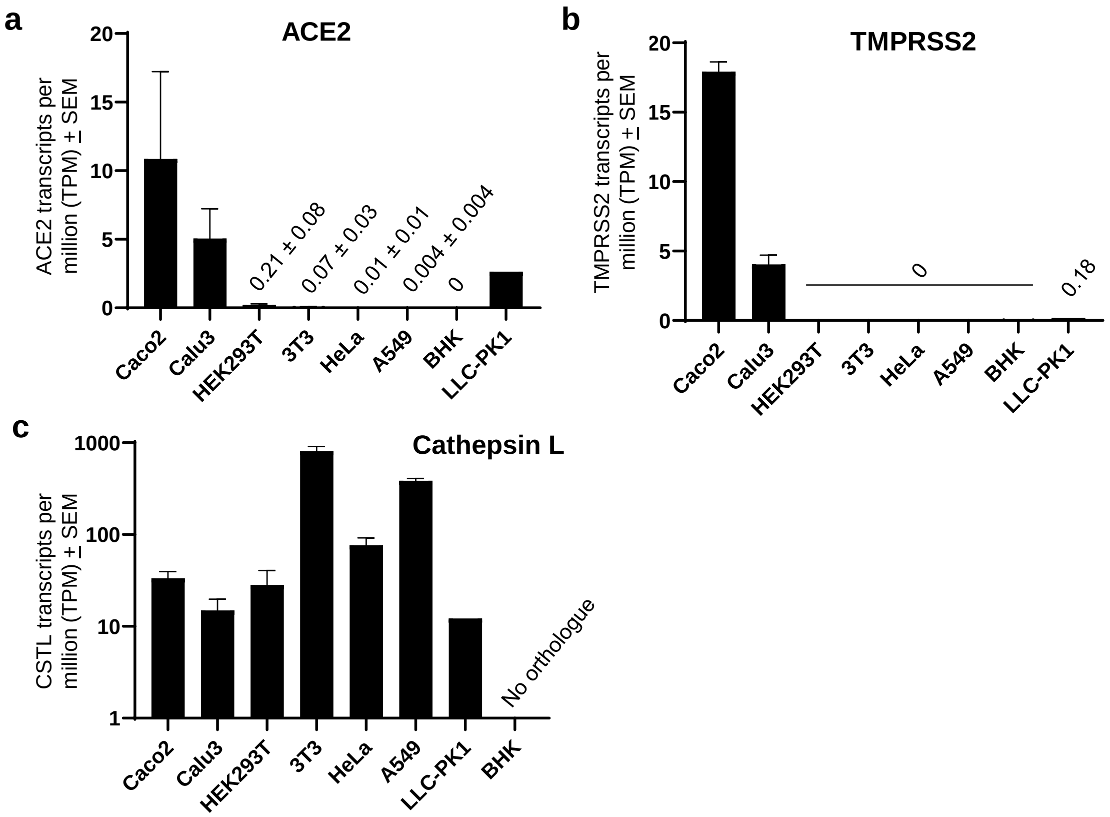


**Supplementary Fig. 5. RNA-Seq read counts for cell lines in Fig. 3.** The cell lines Caco2 and Calu3 can both be infected by SARS-CoV-2 as these express sufficient levels of ACE2[1]. The other cell lines used in Fig. 3 were; 3T3 cells (mouse embryonic fibroblasts), AE17 cells (mouse lung mesothelioma), BHK-21 cells (baby hamster kidney fibroblasts), A549 cells (human lung epithelial carcinoma), HeLa cells (human cervical cancer) and LLC-PK1 cell line (porcine kidney epithelium). SRA run accessions; Caco 2 (n=5), Calu3 (n=4), HEK293T (n=6), 3T3 (n=5), HeLa (n=6), A549 (n=6), BHK (n=3) and LLC-PK1 (n=1). **a** ACE2 transcripts per million reads (TPM) for the indicated cell lines. Although porcine ACE2 is able to support SARS-CoV-2 entry[2] , in LLC-PK1 cells porcine ACE2 is located in the cytoplasm, and not the cell surface (Mori et al, 2022). **b** As for a for TMPRSS2. **c** As for a for Cathepsin L.

[1]Chu et al. 2020. Comparative tropism, replication kinetics, and cell damage profiling of SARS-CoV-2 and SARS-CoV with implications for clinical manifestations, transmissibility, and laboratory studies of COVID-19: an observational study. The Lancet Microbe 1:e14-e23.

[2] Liu et al. 2021. Functional and genetic analysis of viral receptor ACE2 orthologs reveals a broad potential host range of SARS-CoV-2. PNAS 118; e2025373118.

**Method**

Raw data (fastq files) from RNA-Seq experiments were obtained from the Sequence Read Archive (SRA). Fastq files were trimmed of adapter sequences using Cutadapt, mapped to the human reference genome GRCh38 or the mouse reference genome GRCm39 using STAR aligner and TPM normalized gene counts were generated using RSEM. SRA run accessions; HEK293T = SRR16495652, SRR16495651, SRR16495650, SRR16218637, SRR16218638. HeLa = SRR15733492, SRR15733493, SRR15733494, SRR16904840, SRR16904840, SRR16904842. 3T3 = SRR14067064, SRR14067065, SRR9326749, SRR9326749, SRR9326751. A549 = SRR16201279, SRR16201280, SRR16201280, SRR15410446, SRR15410446. Caco2 = SRR13493599, SRR13493596, SRR10416443, SRR10416444, SRR10416444. Calu3 = SRR12709014, SRR12709014, SRR11234095, SRR11234094. BHK-21 = SRR8354394, SRR8354395, SRR8354396. LLC-PK1 = SRR10582029.

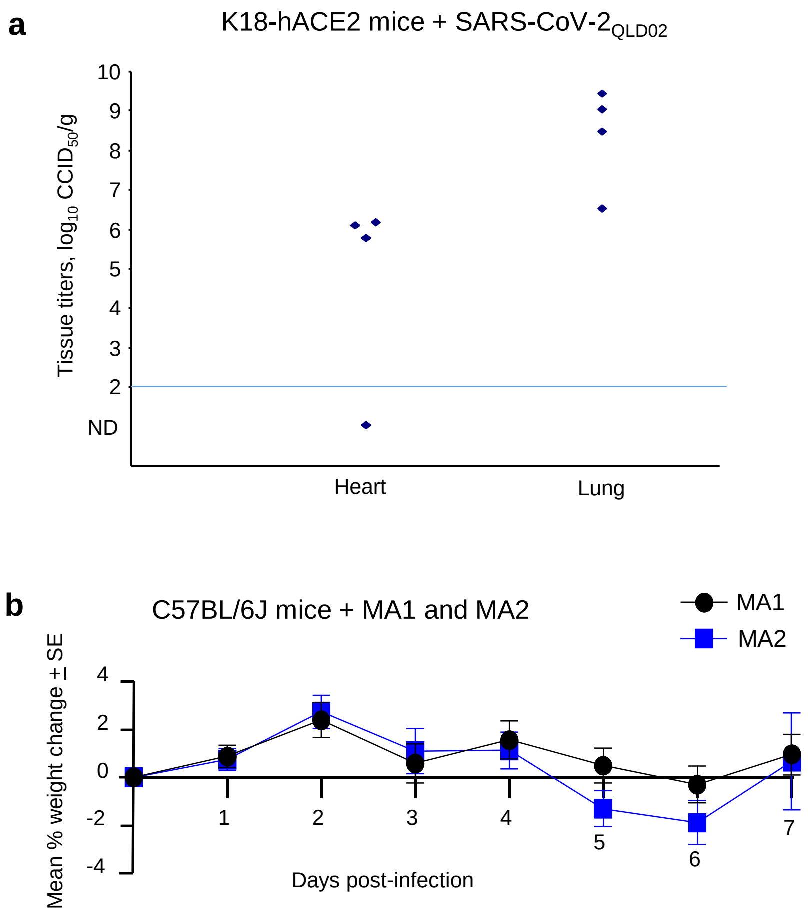


**Supplementary Fig. 6. Heart infection of K18-hACE2 mice and percent weight change for mice in Fig. 4. a** Infection of K18-hACE2 mice with SARS-CoV2$_{QLD02}$ as described (Amarilla et al. 2021). CCID$_{50}$ titers for lung and heart tissues on day 5 post infection (n=4 mice). Heart infection was also reported by Winkler et al. 2020. **b** MA1 and MA2 infection of C57BL/6J mice, percent weight change over time; n=24 on day 2, n=16 on day 4 and n=8 on day 7. Data represents the mean percent weight change relative to day 0.

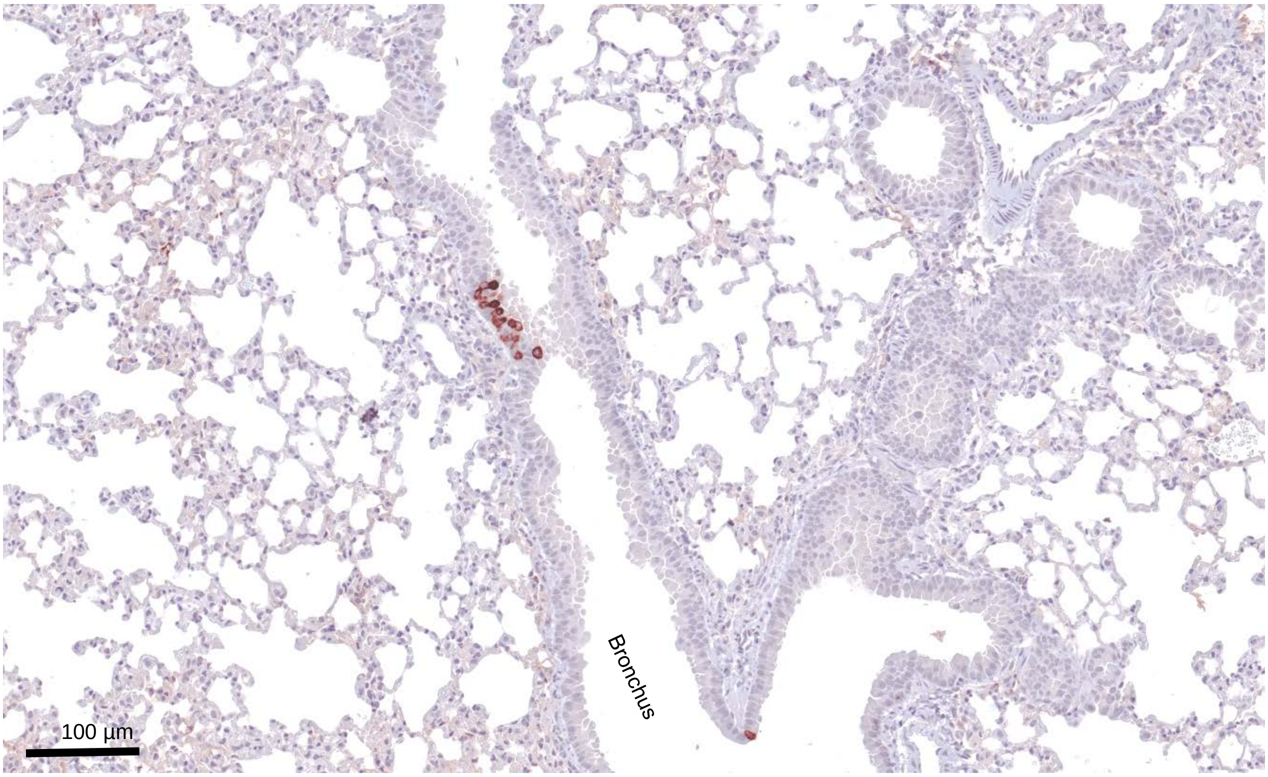


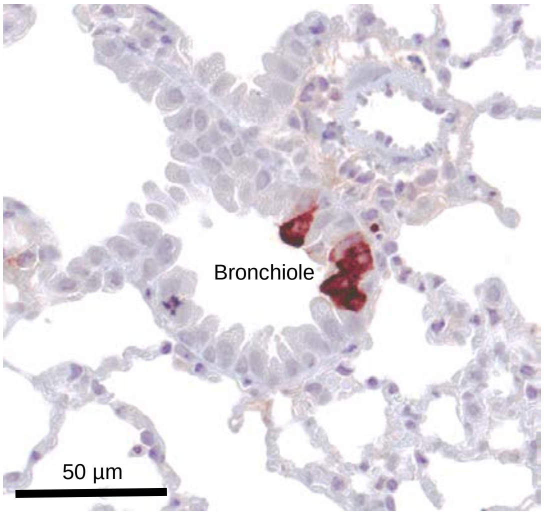


**Supplementary Figure 7. MA2 IHC on day 2 post infection.** IHC using an anti-SARS-CoV-2 spike monoclonal antibody and lungs taken day 2 after infection of C57BL/6J mice with MA2. Images are representative of lung sections from 4 mice. Dark brown staining (Nova Red) indicates infected cells, often with the expected, clearly discernable, cytoplasmic staining pattern. The large unstained areas are included to illustrate the specificity of the IHC staining.

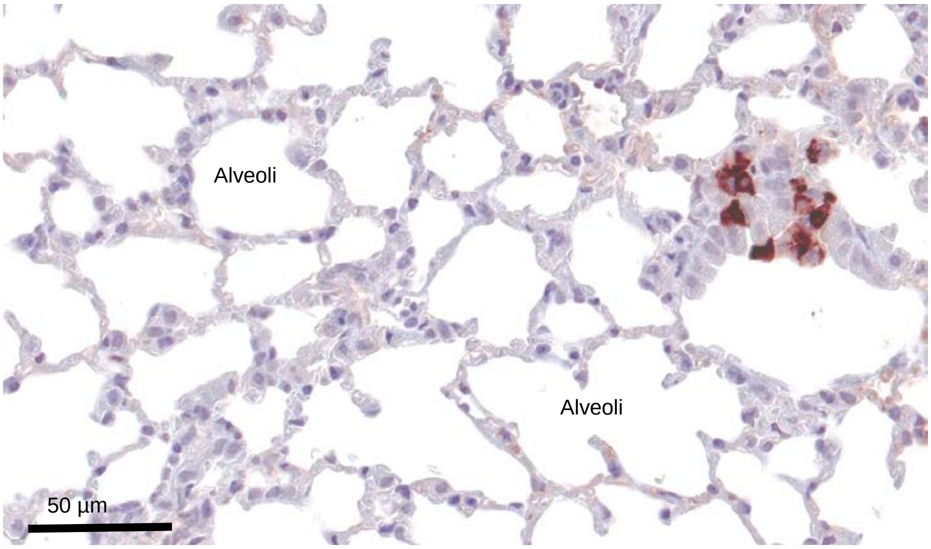

**a** Occasional prodigious staining of bronchial epithelium

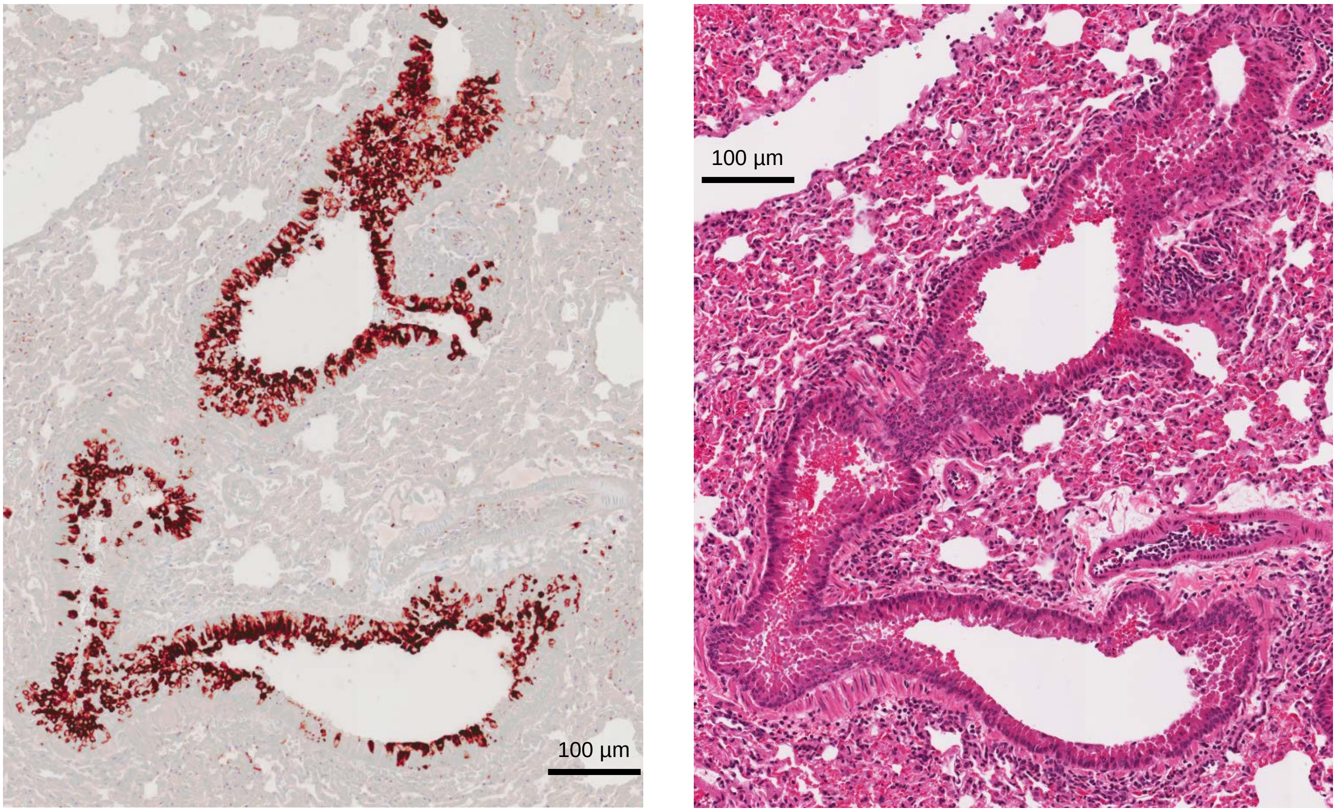


**b** Occasional staining of columnar epithelial cells in the trachea

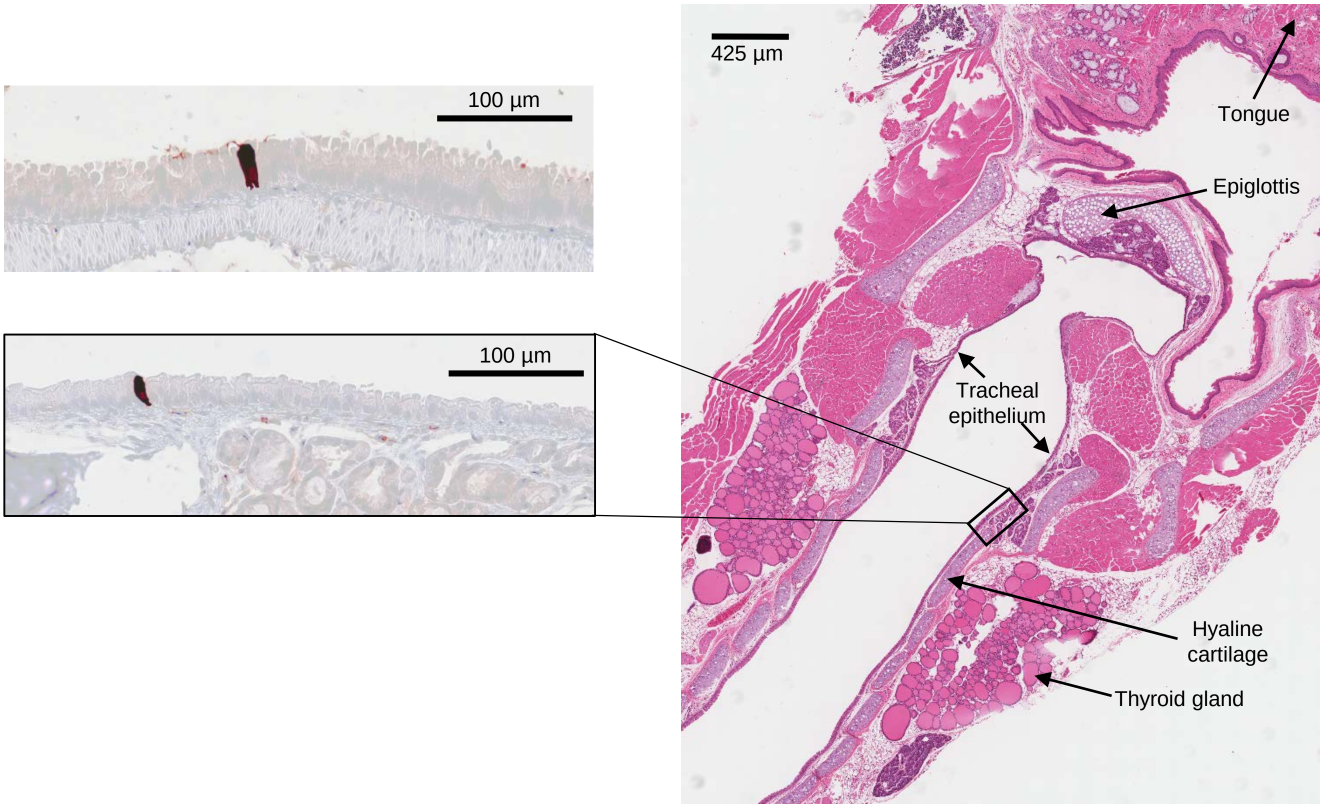


**Supplementary Fig. 8. Additional lung and trachea IHC and H&E for MA1. a** C57BL/6J mouse lungs harvested at day 2 post-infection (MA1) underwent anti-spike IHC as per Figure 4d-g. An example of prodigious bronchial epithelial staining is shown (left), alongside the corresponding H&E section (right). **b** Anti-spike IHC staining showing occasional positive columnar epithelial cells in the trachea (left). H&E (right) provides anatomical location for the inset IHC image.

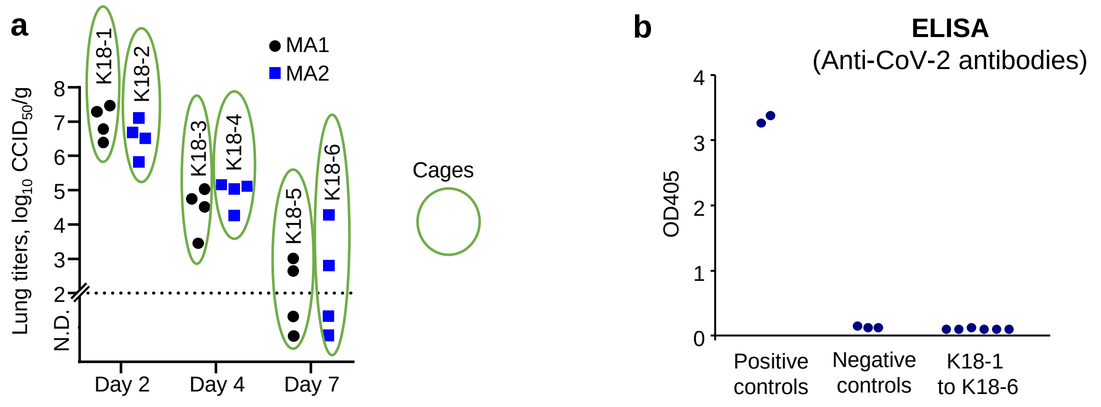


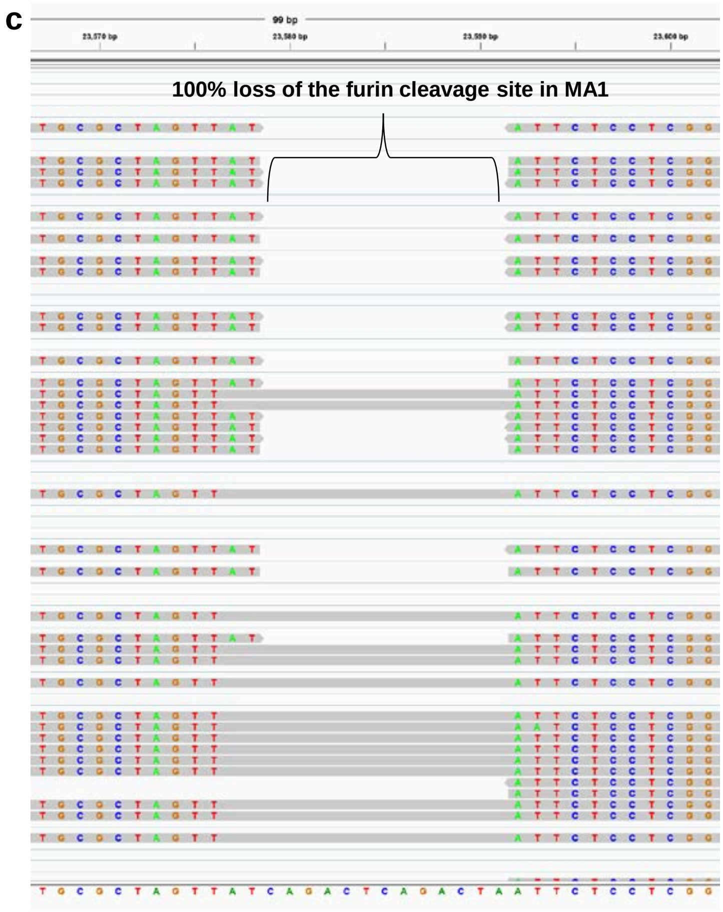


**Supplementary Fig. 9. No evidence of mouse to mouse transmission for MA1 or MA2.** **a** Mice shown in Fig. 5A were housed in 6 cages (green ovals) and contained MA-infected C57BL/6J mice that were culled on day 2, 4 or 7 post infection. Fig. 5A is reproduced here to show cage set up. Each cage also contained one uninfected K18-hACE2 mouse (K18-1 through to K18-6). **b** Three weeks after infection of the C57BL/6J mice with MA1 or MA2, the K18-1 to 6 mice were bled and tested for anti-CoV-2 antibodies as a indicator of infection (as described Bao et al. 2020). Positive controls K18-hACE2 mice were immunised with UV-inactivated virus, with serum tested in duplicate at a 1 in 150 dilution (n=2 mice). Negative controls - naive mice, tested in duplicate at a 1 in 10 dilution of serum (n=3 mice). K18-1 to K18-6 – the six mice co-caged with infected mice shown in a, sera tested in duplicate at 1 in 10 dilution. ELISA antigen was UV-inactivated QLD02. **c** RNA-Seq of MA1 provided >40,000 reads flanking the furin cleavage site. IGV of a subset of reads is shown. No reads were found that showed an intact furin cleavage site. The furin cleavage site has been identified as critical for transmission in ferrets and is a well described virulence determinant in humans. This deletion therefore provides a built-in safety feature for MA1, with a loss of around 13 nucleotides not likely to be readily reversible in a laboratory setting.

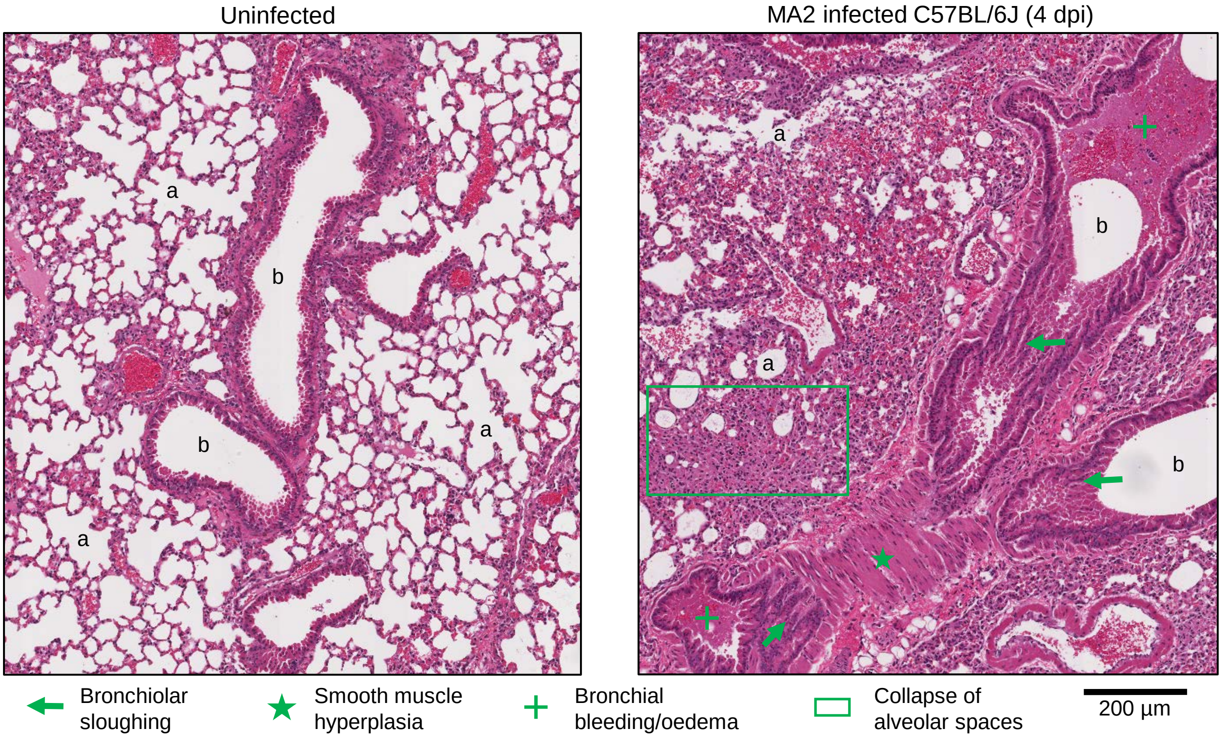


**Supplementary Figure 10. H&E staining of lungs after infection of C57BL/6J mice with MA2.** Representative low resolution images of H&E stained lung sections for uninfected (left) and MA2 infected (right) C57BL/6J mice taken 4 days post infection. Examples of bronchiolar sloughing (green arrow), smooth muscle hyperplasia (green star), bronchial bleeding/oedema (green plus sign), and collapse of alveolar spaces (green rectangle), are shown.

**a** E484D (RBD)

| Position 484 | No. GSIAD |
|---|---|
| A | 1,832,377 |
| V | 1423 |
| L | 13 |
| I | 8 |
| P | 50 |
| M | 15 |
| F | 1 |
| W | 1 |
| G | 374 |
| S | 97 |
| T | 51 |
| C | 3 |
| Y | 0 |
| N | 4 |
| Q | 22581 |
| **D** | **298** |
| E | QLD02 |
| K | 254,598 |
| R | 44 |
| H | 2 |

**b**

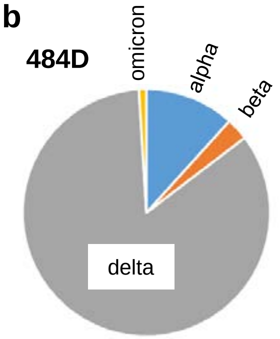


**c**

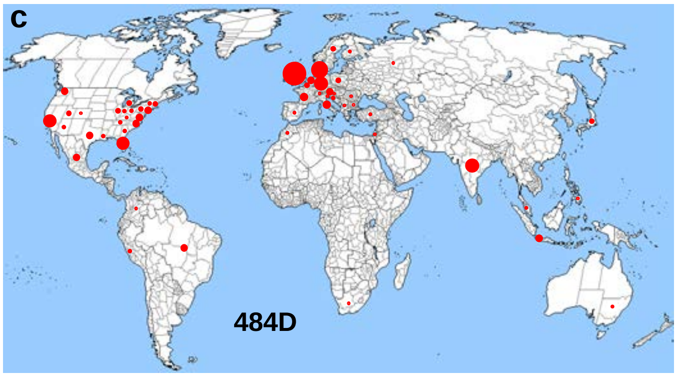


**d** Q498H (RBD)

| Position 498 | No. GSAID |
|---|---|
| A | 2 |
| V | 3 |
| L | 79 |
| I | 3 |
| P | 21 |
| M | 24 |
| F | 4 |
| W | 0 |
| G | 0 |
| S | 12 |
| T | 10 |
| C | 1 |
| Y | 16 |
| N | 20 |
| Q | QLD02 |
| D | 10 |
| E | 31 |
| K | 48 |
| R | 1754779 |
| **H** | **77** |

+ve charged aa

**e** P812L

| Position 812 | No. GSAID |
|---|---|
| A | 51 |
| V | 3 |
| **L** | **9085** |
| I | 3 |
| P | QLD02 |
| M | 2 |
| F | 9 |
| W | 0 |
| G | 0 |
| S | 10003 |
| T | 330 |
| C | 2 |
| Y | 0 |
| N | 5 |
| Q | 150 |
| D | 1 |
| E | 0 |
| K | 4 |
| R | 272 |
| H | 1 |

**f** P812

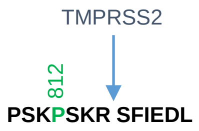


**g** E96A

| Position 96 | No. GSIAD |
|---|---|
| **A** | **253** |
| V | 1526 |
| L | 88 |
| I | 0 |
| P | 156 |
| M | 1 |
| F | 5 |
| W | 2 |
| G | 663 |
| S | 179 |
| T | 5 |
| C | 1 |
| Y | 27 |
| N | 13 |
| Q | 1451 |
| D | 2253 |
| E | QLD02 |
| K | 41 |
| R | 21 |
| H | 6 |

**h** D215G

417N
484K
501Y

RBD

18F
80A
246I
701V
**215G**

Lineage defining amino acids in Spike of beta variants

**i** RNA Seq of QLD02 stock virus

| | Read count | A | C | G | T | N | QLD02 | | MA1 | |
|---|---|---|---|---|---|---|---|---|---|---|
| E96A | 92059 | 91847 | 69 | 111 | 27 | 5 | E | GAG | A | GCG |
| D215G | 90528 | 80020 | 146 | 10340 | 19 | 3 | D | GAG | G | GGG |
| E484D | 135035 | 134978 | 5 | 16 | 32 | 4 | E | GAA | D | GAC |
| Q498H | 173155 | 172877 | 93 | 101 | 75 | 9 | Q | CAA | H | CAC |
| P812L | 181899 | 194 | 181632 | 20 | 45 | 8 | P | CCA | L | CTA |

**Supplementary Fig. 11. MA1 spike substitutions.** **a** D in position 484 (484D) of the receptor binding domain (RBD) has been associated with ACE2-independent infection and 298 human virus isolates submitted to GISIAD have D in this position. Other amino acids, such as A (in omicron, yellow), as well as (most commonly) V, G, Q and K are also found. **b** Viruses with 484D are found in all variants of concern, but mostly in delta. **c** Viruses with 484D are found globally. Circle size is to scale for the number of submissions with 484D in GISAID (total = 298). This is raw data; no correction for number of submissions per country. **d** Viruses with H in position 498 are present in 77 human isolates in GISAID. This site contains R in omicron strains (yellow). H and R both contain positively charged side groups. **e** Viruses with L in position 812 are present in 9085 human isolates in GISAID. **f** Position 812 is located near the S2′ cleavage site; cleaved by TMPRSS2 (blue arrow). **g** Viruses with A in position 96 are present in 253 human isolates in GISAID. **h** G in position 215 is one of eight lineage defining amino acids in Spike for beta variants of concern. **i** RNA-Seq of QLD02 stock showing that MA1 substitutions (green) might be present in QLD02 stock; however, only 215G (11.4% of QLD02 stock) was clearly higher than the sequencing error rate (Q30 ≈ 1 in 1000). Dominant nucleotides in QLD02 – blue numbers and text.

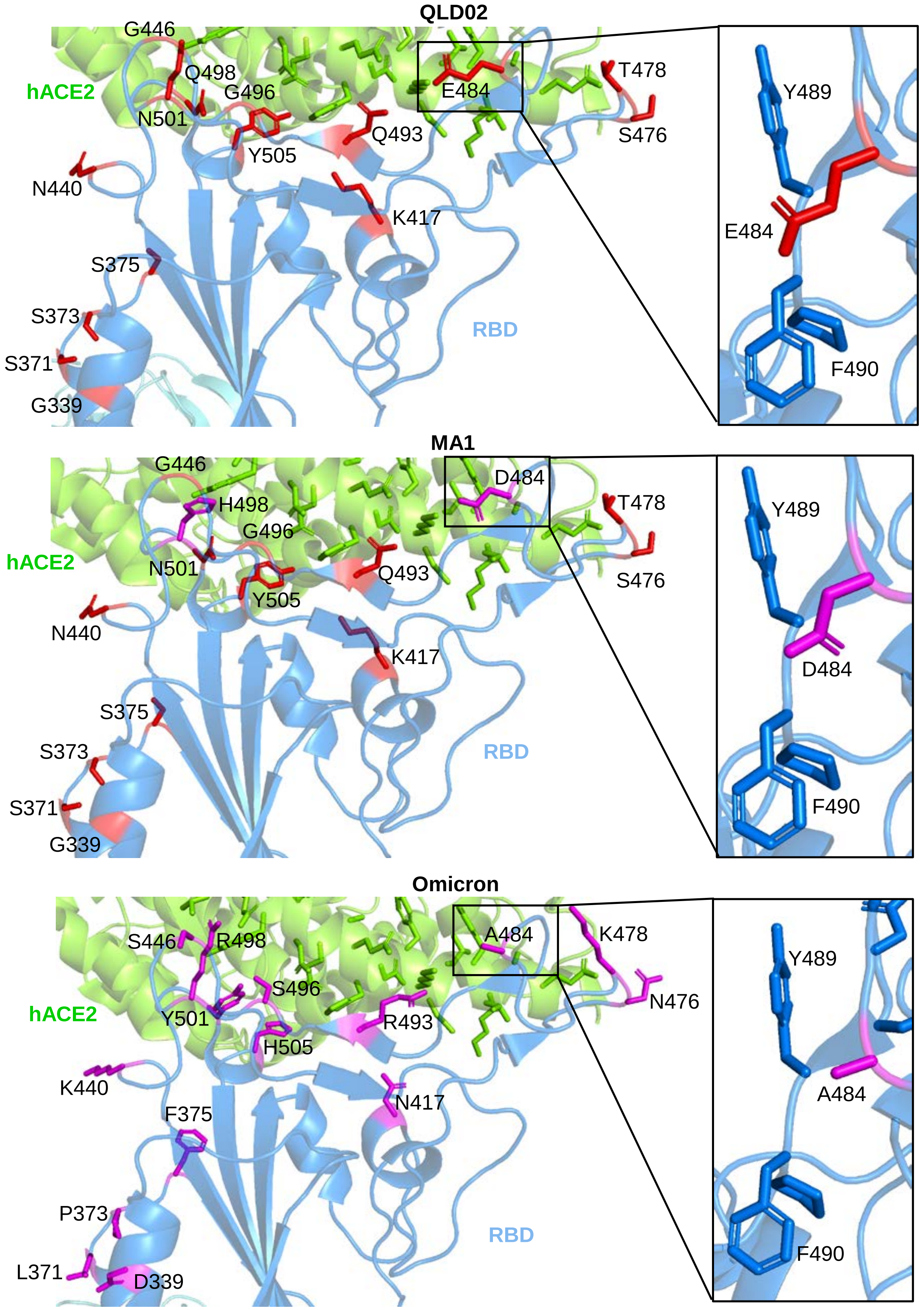


**Supplementary Fig. 12 RBD amino acid changes in MA1 and the Omicron variant.** The structure of the spike RBD bound to hACE2 (PDB: 7df4) viewed in PyMOL. Red amino acids = QLD02 residues, and purple amino acids = changes from QLD02 in MA1 or Omicron variant. Inset is zoomed in on position 484.

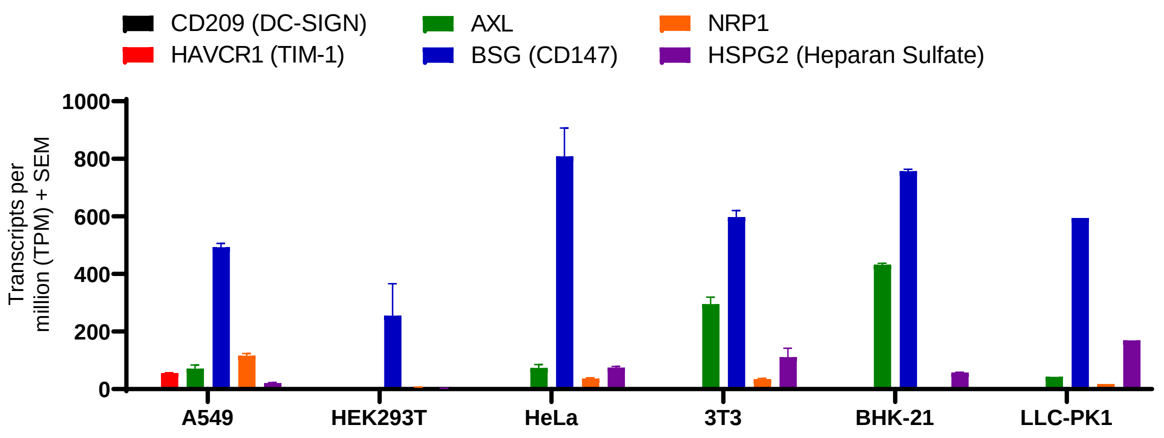


**Supplementary Fig. 13. mRNA expression for putative alternative receptors in the cell lines tested in Fig. 3.** Mean + SEM transcripts per million (TPM) for DC-SIGN (black), TIM-1 (red), AXL (green), CD147 (blue), NRP1 (orange), and HSPG2 (purple), in the following cell lines; A549 (n=6), HEK293T (n=6), HeLa (n=6), 3T3 (n=5), BHK-21 (n=3) and LLC-PK1 (n=1). Publically available data was downloaded from Sequence Read Archive (SRA) (listed in Supplementary Fig. 5).
